# CRISPR/Cas9 deletions induce adverse on-target genomic effects leading to functional DNA in human cells

**DOI:** 10.1101/2021.07.01.450727

**Authors:** Keyi Geng, Lara G. Merino, Linda Wedemann, Aniek Martens, Małgorzata Sobota, Yerma P. Sanchez, Jonas Nørskov Søndergaard, Robert J. White, Claudia Kutter

## Abstract

The CRISPR/Cas9 system is widely used to permanently delete genomic regions by inducing double-strand breaks via dual guide RNAs. However, on-target consequences of Cas9 deletion events have yet to be fully investigated. To characterize Cas9-induced genotypic abnormalities in human cells, we utilized an innovative droplet-based target enrichment approach followed by long-read sequencing and coupled it to a customized *de novo* sequence assembly. This approach enabled us to dissect the sequence content at kilobase scale within an on-target genomic locus. We here describe extensive genomic disruptions by Cas9, involving a genomic duplication and inversion of the target region as well as integrations of exogenous DNA and interchromosomal DNA fragment rearrangements at the double-strand break sites often at the same time. Although these events altered the genomic composition of the on-target region, we found that the aberrant DNA fragments are still functional, marked by active histones and bound by RNA polymerase III. In HAP1 cells, the integration of the target-derived fragments accelerated cell proliferation in deletion clones. Our findings broaden the consequential spectrum of the Cas9 deletion system, reinforce the necessity of meticulous genomic validations and rationalize extra caution when interpreting results from a deletion event.

## INTRODUCTION

The CRISPR/Cas9 system has revolutionized genome engineering approaches. Various toolsets have been developed, which allow loss-of-function perturbations of functional genomic elements in a simple and efficient manner. To modify the genome, the components of the CRISPR/Cas9 system are exogenously introduced into human cells (1–3). In the presence of a guide RNA (gRNA) that is complementary to the target site and next to a protospacer adjacent motif (PAM), the expressed Cas9 endonuclease induces a double-strand break (DSB) at a specific genomic site. If an exogenous homologous DNA template is supplied, the DSB is repaired by the homology-directed repair to introduce specific mutations or insertions of desired sequences (4). Alternatively, microhomology-mediated end joining (MMEJ) can be triggered by short (5 to 25 bp) overlapping sequences that allow for recombination at the DSB (5). In contrast, in the absence of a homologous template, the cell uses non-homologous end joining (NHEJ) to resolve the DSB. This usually results in small deletions or insertions of a few nucleotides (1). Instead of introducing single nucleotide or short sequence deletions, longer DNA regions can be excised from the genome by using dual gRNAs that flank the target region and guide Cas9 to induce two DSBs (6, 7). Through this approach, a target region of about 1 Mb could successfully be removed from the genome in mouse cells (8). In addition, a variety of genomic regions have been modified using the dual gRNA system. For example, the genomic manipulation of enhancers (9, 10), chromatin loop anchors (11, 12), protein-coding genes (7, 13) and noncoding genes (14) revealed disease-associated functional elements and genes. Although a broad range of functional sequences has been successfully deleted from the genome, transfer RNA (tRNA) genes have not been targeted by using the dual gRNA system in human cells.

Nuclear encoded tRNA genes are transcribed by RNA polymerase III (Pol III) through the recognition of internal promoter binding motifs (15, 16). Pol III occupancy of tRNA genes is widely used to quantify tRNA gene usage (17–24). In addition, Pol III-bound tRNA genes reside in euchromatic genomic regions, which are marked by active histones, such as histone 3 lysine 4 trimethylation (H3K4me3) (15, 21, 22, 25, 26). tRNAs represent one of the most abundant RNA types. These circa 73 nt long RNA molecules act as physical adapter molecules during translation, in which the anticodon region of a tRNA molecule recognizes the complementary codon of a messenger RNA and adds an amino acid to the growing polypeptide chain. tRNAs have a wide spectrum of additional functions. For example, a tRNA molecule can be cleaved into tRNA halves or fragments, which impair gene regulation transcriptionally or post-transcriptionally and contribute to diseases, such as cancers and neurodegenerative disorders (16, 27–32). Furthermore, in some instances, the tRNA gene locus itself functions in genome organization by acting as an insulator (33, 34).

In order to investigate tRNA gene usage, we deleted two tRNA genes from the genomes of human near-haploid chronic myeloid leukemia (HAP1) and hyperploid hepatocellular carcinoma (HepG2) cells using the CRISPR/Cas9 system with dual gRNAs. By applying an advanced droplet-based target enrichment method (Xdrop) (35) followed by Oxford Nanopore Technology (ONT) long-read sequencing (LRS) (36) enabled us to uncover several unexpected genomic alterations at the target site. Although Cas9 induced genomic cuts, and our target region was successfully excised, we found complex genomic aberrations, including duplication and inversion of the target region, as well as integration of exogenous and interchromosomal DNA fragments. Despite our initial attempt to delete tRNA genes to impair their expression, the aberrant genomic modifications at the original locus resulted in actively transcribed tRNA genes in HAP1 and HepG2 cells. This highlights that CRISPR/Cas9-based genomic engineering can cause undesired on-target effects. Our work also underscores the complexity of human DNA repair mechanisms in the presence of the powerful prokaryotic Cas9 nuclease and raises serious concerns when studying the functionality of modified genomes. We therefore recommend examining modified cells to a greater extent through long-read sequencing of the on-target locus since several unintended genomic alterations may occur at the same time and could be missed by standard Sanger sequencing and PCR verification methods.

## RESULTS

### The target region remained detectable and functional in Cas9 deletion clones

To test the effectiveness of CRISPR/Cas9 for deleting tRNA genes, we focused on a pair that is closely located on the human chromosome 17 (**Fig. 1A**). Since tRNA genes belong to one of the largest multi-copy gene families (25), in which individual gene family members are identical in sequence composition, we designed gRNAs mapping to the unique 5’ and 3’ flanking regions. That enabled Cas9 to excise an 870 bp long genomic fragment containing the two tRNA genes (Δt) (**Fig. 1B**). We transfected HAP1 and HepG2 cells with two Cas9 plasmids carrying one of the two gRNAs and a small size plasmid to enhance transfection efficiency (Material and Methods, **Fig. S1A**). Since the CRISPR/Cas9 vector contains the puromycin resistance gene, we used antibiotic selection to identify positively transfected cells and propagated further single cell-derived clones. To validate the clonal deletion of the target region, we performed a PCR with flanking region-specific primers. The size of the PCR product was indicative of a successful deletion when inspected by agarose gel electrophoresis and its sequence content was verified by Sanger sequencing (**Fig. 1B, Fig. S1A**). Overall, 94 HAP1 and 90 HepG2 single cell-derived clones were generated. Among them, 5 HAP1 and 17 HepG2 clones contained a deletion (**Fig. S1B-D**). We focused our subsequent analysis on two validated deletion clones, referred to as HAP1 Δt72 and HepG2 Δt15 (**Fig. S1B-D**).

**Fig 1.**
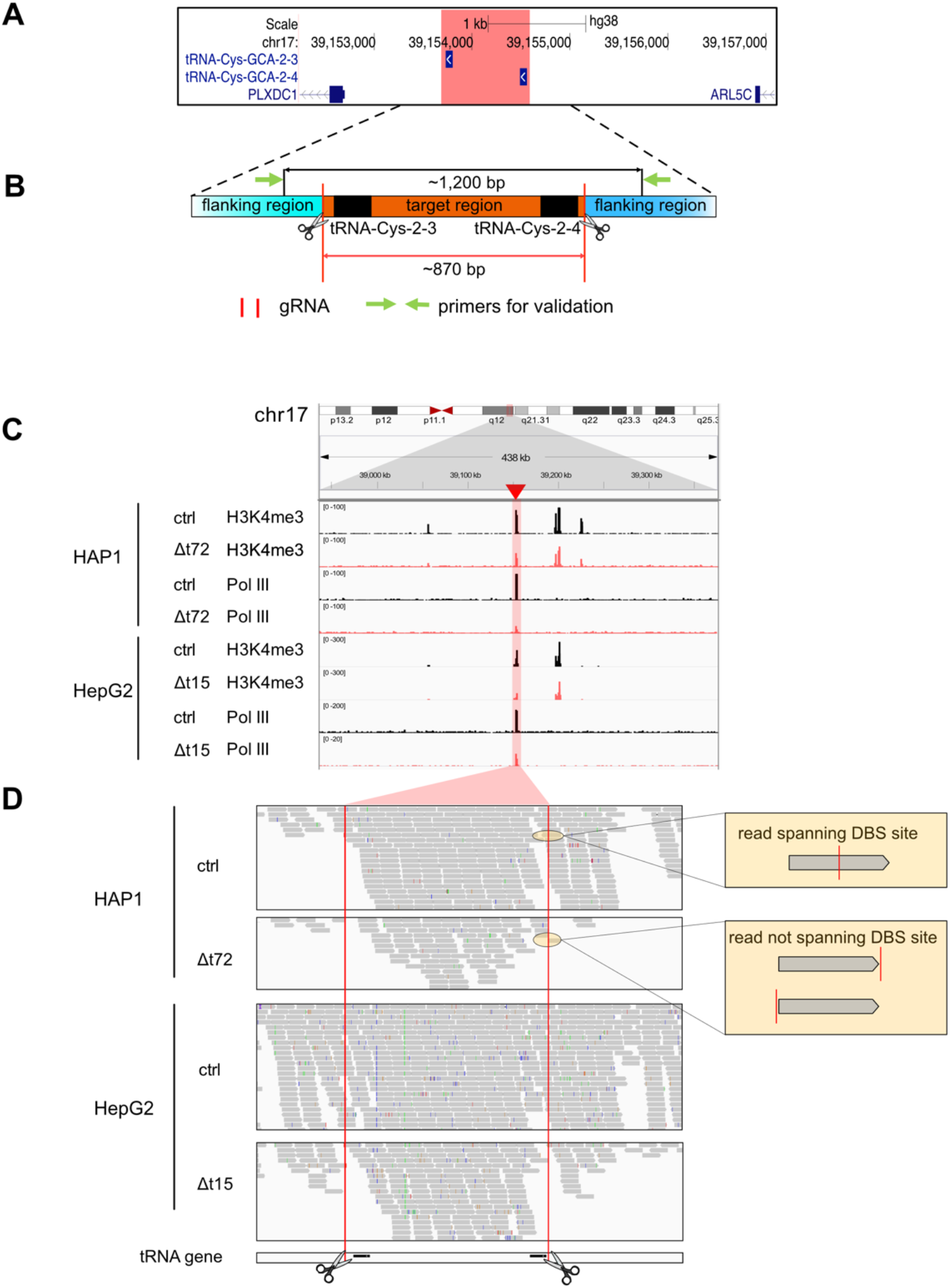
The target region remained functional in deletion clones. **(A)** The hg38 genome browser view shows the genomic location of our target locus (red). Arrows denote directionality of gene transcription. **(B)** Illustration of the design strategy of Cas9 dual gRNA deletion strategy. Cas9 cut sites (red horizontal lines and scissors) and primers for validation (green arrows) are indicated. The target region (orange) containing two target tRNA genes (black) is around 870 bp. The size of the PCR product in control clones is around 1,200 bp. **(C)** The hg38 genome browser view demonstrates normalized ChIP-seq reads for H3K4me3 and Pol III covering the target loci (highlighted in red) in the HAP1 control (ctrl) and Δt72 as well as HepG2 ctrl and Δt15 clones. **(D)** Alignment tracks show individual H3K4me3 ChIP-seq reads of the deleted and surrounding region as in (C). DSB sites (red lines and scissors) and tRNA gene locations (black) are indicated. Examples of reads spanning (top) or not spanning (bottom) DSB site are illustrated (yellow box).

To study the consequences of tRNA gene deletions, we utilized published Pol III ChIP-seq data in HepG2 unmodified cells (23) and profiled the genome-wide binding of H3K4me3 and Pol III in both deletion and non-targeting control (ctrl) clones (**Fig. 1C**). We only considered reads with high mapping quality (Material and Methods). Surprisingly, we observed binding of H3K4me3 and Pol III to DNA sequences within the target region in HAP1 and HepG2 deletion clones, although at a lower magnitude, when compared to their respective control clones (**Fig. 1C**). To explain this observation, we considered several possibilities that can lead to imperfect genomic Cas9 deletions in cells. Previous reports described genomic aberrations when using the Cas9 dual gRNA system. For example, (I) the application could lead to heterozygous deletion clones (8) whereby one allele carries the deletion and the other allele is unmodified, (II) mutations could occur at the PAM sequences (54, 55) preventing the recruitment of Cas9 and its cutting activity at the gene locus, (III) the Cas9 cleavage might be unsynchronized at the two DSB sites (56), in which one DSB would have been repaired before the induction of the second DSB or the target sequence could have been excised and then either (IV) inverted or (V) duplicated (8, 13, 57–59). However, to our knowledge a combination of a simultaneously occurring inversion and duplication has not been described. Scenarios I to IV can be excluded since we would have detected a PCR product corresponding to the size of the unmodified allele (circa 1,200 bp) (**Fig. S1C-D**). We also excluded scenario V since we found ChIP-seq reads spanning the excision site in the control but not in the deletion clones (**Fig. 1D**). This prompted us to consider alternative consequences occurring as a result of Cas9 genome engineering.

### Applying Xdrop in deletion clones confirmed the genomic remodeling of the target region

The Xdrop technology has recently been applied to validate CRISPR/Cas9 genome modifications (35, 48). To investigate the on-target editing outcomes in our Cas9 deletion clones, we applied the Xdrop approach to enrich for sequences containing our CRISPR/Cas9-targeted genomic region in the HAP1 Δt72 and HepG2 Δt15 deletion clones. Subsequently, we identified the sequence content of the Xdrop-enriched molecules by ONT LRS (Xdrop-LRS). On average, we obtained 217,000, and 179,000 reads of a median size of 4,600 and 5,200 bp in the HAP1 Δt72 and HepG2 Δt15 deletion clones, respectively. Read coverage at the target locus in each cell clone indicated sufficient enrichment (**Fig. S2A-B**). By aligning both corrected and raw reads to the human reference genome, we observed sharp declines in coverage at the two DSB sites and no read spanned the two DSB sites in these two deletion clones (**Fig. S2A**). This supported our conclusion drawn from our ChIP-seq data that the target region was altered in the HAP1 Δt72 and HepG2 Δt15 deletion clones.

### A *de novo* assembly-based approach revealed a duplication, inversion and local insertion of the target region in the HAP1 deletion clone Δt72

To decipher the underlying alterations that caused a rearrangement of the CRISPR/Cas9-targeted region, we employed a customized *de novo* sequence assembly approach (**Fig. S2C**). For this, we assembled contigs containing the target region and connected them with sequences of the respective flanking regions for each of the two deletion clones.

In our HAP1 Δt72 deletion clone, we identified contigs with three distinct breakpoints that deviated from the human reference genome composition (**Fig. 2A**). One breakpoint (BP2) connected two units of our deleted target region in tandem orientation. This finding is indicative of a duplication event of our original target fragment. The other two breakpoints (BP1 and BP3) connected the duplicated fragments with the 5’ and 3’ flanking region of our original target region in inverse orientation. This suggested an unexpected event in which the deleted fragment of our target region got duplicated, inverted and inserted at the original gene locus (**Fig. 2A**). To eliminate any confounding effects from our approach, we inspected both raw and corrected read alignment tracks. We found multiple long reads spanning the contig and several additional reads covering the individual breakpoints confirming on-target genomic alteration events (**Fig. 2A**).

**Fig 2.**
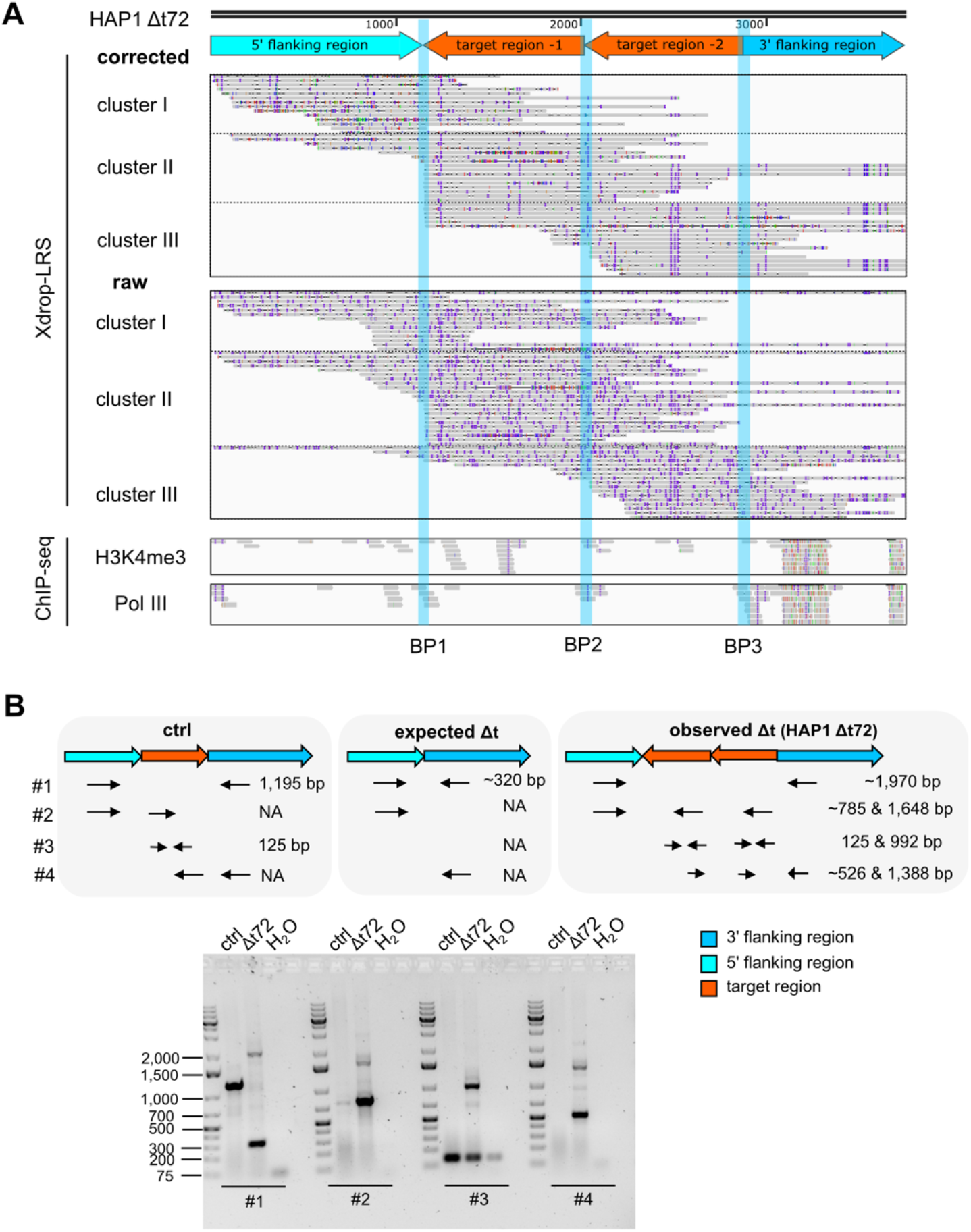
A duplication, inversion and local insertion of target-derived fragments occurred in the HAP1 Δt72 deletion clone. **(A)** The hg38 genome browser view shows Xdrop-LRS (top, corrected and raw reads) and ChIP-seq (bottom, H3K4me3 and Pol III) reads aligned against the assembled contig. Arrows (top) show genomic orientation and approximate size of the flanking (5’ light and 3’ dark blue) and target (orange) regions within the contig. Representative aligned reads for cluster I, II and III support three different breakpoints (BP1-3, blue vertical lines) within the contig. **(B)** Schematic illustration of the primer design strategy to validate contigs by PCR and the expected amplicon length (top) representing scenarios in an unmodified control, expected and observed deletion event. Agarose gel (bottom) confirms the size of the obtained PCR products. Marker bands specify DNA size in bp.

Since it has been shown that the multiple displacement amplification in droplets (dMDA) can lead to false positive calls of duplication and inversion events (60), we aligned our Illumina ChIP-seq data to the assembled contig. Without allowing soft-clipped reads, we still found multiple H3K4me3 and Pol III ChIP-seq reads mapping to the three BPs and thereby confirmed our detected duplication and inversion events (**Fig. 2A**). In addition, we performed an independent validation of our *de novo* sequence assembly by designing three sets of primers specific for the flanking and/or target region (**Fig. 2B, Table S1**). The different primer combinations and size of PCR products enabled us to discern the genomic composition of our target locus in our unmodified, expected and actually observed deletion clones (**Fig. 2B**). This PCR-based result supported our Xdrop-LRS findings and our conclusion that the target region got duplicated, inverted and inserted in the HAP1 Δt72 deletion clone.

### The target sequence got duplicated, inverted and co-integrated with exogenous DNA fragments in the HepG2 deletion clone Δt15

Similar to the HAP1 Δt72 deletion clone, we employed our customized *de novo* sequence assembly approach to investigate potential genomic aberrations of the target region in the HepG2 Δt15 deletion clone. The assembly of our Xdrop-LRS reads revealed four breakpoints (**Fig. 3A**). We found that a duplication of our target region occurred in the HepG2 Δt15 deletion clone. Breakpoint 2 (BP2) connected these two units of the target sequence but in divergent orientation. This suggested that only one of the two units got inverted, which was further confirmed by BP1 which linked the 5’ flanking region of the target to the duplicated target region itself. Resolving the events at the 3’ cut site revealed that exogenous DNA fragments were integrated at the DSB site. We searched for high sequence similarity of these fragments (Material and Methods) and found that an approximately 200 bp long fragment aligned perfectly to a part of the ribonuclease R (rnr) gene in the *E. coli* genome (**Fig. 3A**). BP3 joined this 200 bp *E. coli* genome fragment and the duplicated target region. In addition, we observed a more than 6,000 bp long fragment mapping to the CRISPR/Cas9 vector that we used for cellular transfections. BP4 connected the *E. coli* fragment to the CRISPR/Cas9 vector sequence. Reads mapping to the CRISPR/Cas9 vector included sequences of the U6 promoter, the gRNA cloning site, the gRNA scaffold, the chicken β-actin promoter and the first half of the Cas9 gene (**Fig. 3B**). Remarkably, we found reads carrying the sequence of gRNA-Δt-1, which we synthesized to induce the DSB at the 5’ flanking region. However, there were no reads containing the sequence of gRNA-Δt-2, suggesting that the gRNA and the scaffold part of the CRISPR/Cas9-gRNA-Δt-1 but not -2 vector got integrated. At a lower coverage, we also noted reads mapping to sequences of the poly(A) signal, the f1 origin of replication (f1 ori), the ampicillin resistance gene and the second ori (**Fig. 3B**). In addition to the raw and corrected Xdrop-LRS reads, we found that H3K4me3 and Pol III ChIP-seq reads mapped to the BPs. Furthermore, our PCR results using primers annealing to different components of the contig supported an accurate contig assembly (**Fig. 3A and C**). Surprisingly, the integrated exogenous DNA fragments originating from the *E. coli* genome and the CRISPR/Cas9 vector were marked by H3K4me3, which indicated active sequence usage (**Fig. 3A**).

**Fig 3.**
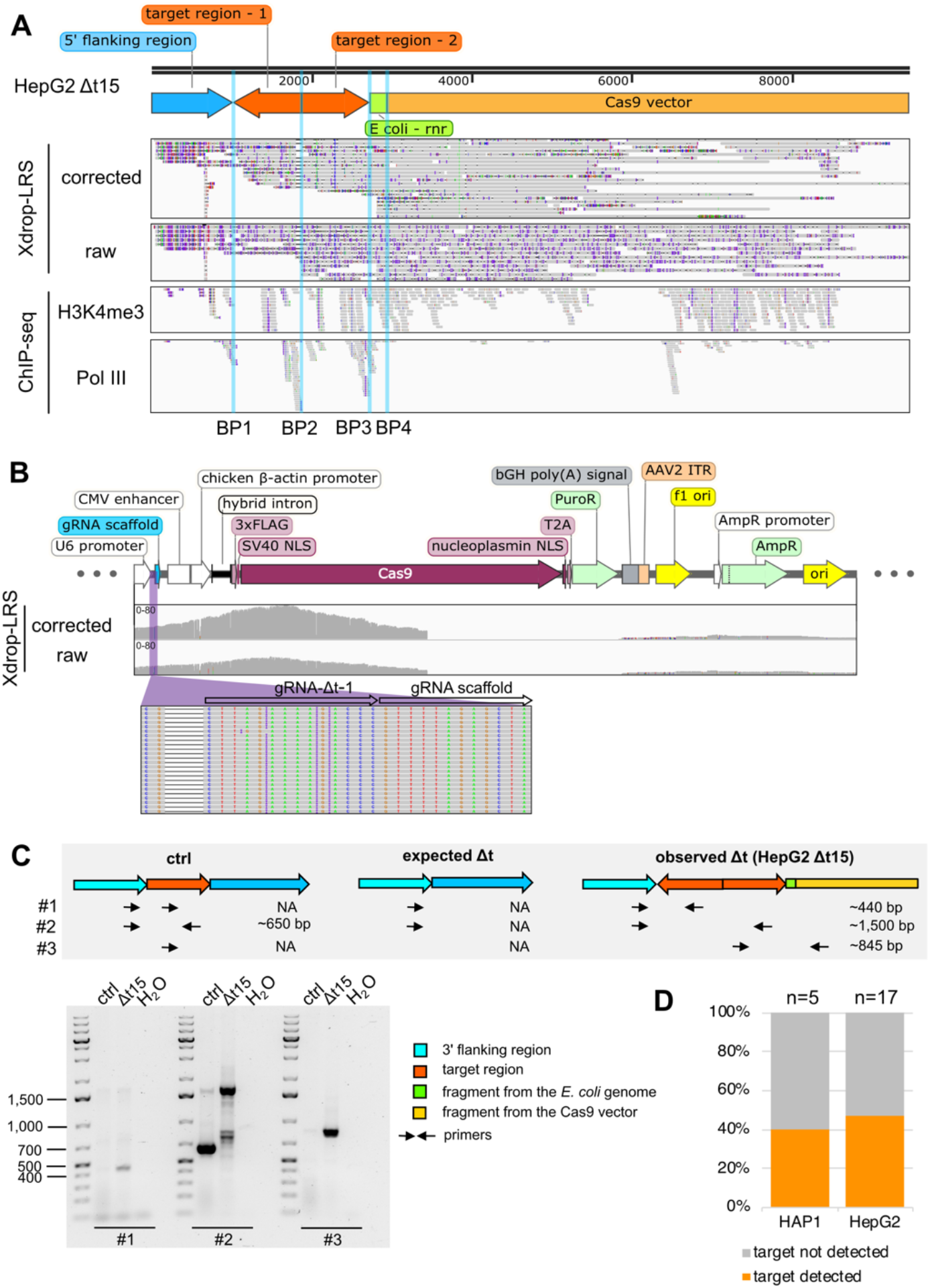
A duplication, inversion of target-derived fragments and integration of exogenous DNA fragment arose in the HepG2 Δt15 deletion clone. **(A)** The hg38 genome browser view shows Xdrop-LRS (top, corrected and raw reads) and ChIP-seq (bottom, H3K4me3 and Pol III) reads aligned against the assembled contig supporting four different breakpoints (BP1-4, blue vertical lines) within the contig. Arrows (top) show genomic orientation and approximate size of the flanking (blue) and target (orange) regions as well as sequences of the *E. coli* genome (green) and the CRISPR/Cas9-gRNA-Δt-1 transfection vector (yellow) present within the contig. **(B)** Coverage tracks display Xdrop-LRS corrected and raw reads aligning to the CRISPR/Cas9 vector (top). Sequence annotations and directions of gene transcription are labelled in accordance to the CRISPR/Cas9 vector map. The gRNA-Δt-1 cloning site and reads mapping to this region are highlighted (purple horizontal line) and magnified (bottom). **(C)** Schematic illustration of the strategy to validate the contig and the expected PCR product length (top) in an unmodified control, expected and the HepG2 Δt15 deletion clone. Agarose gel electrophoresis (bottom) visualizes the size of the PCR products. Marker bands specify DNA size in bp. **(D)** Stacked bar plot indicates the frequency of on-target genomic alterations in the validated HAP1 (n=5) and HepG2 (n=17) Δt deletion clones.

Since we co-transfected the CRISPR/Cas9 and the pBlueScript vector to increase cell transfection efficiency (3), we inspected whether sequences of the pBlueScript vector could have integrated. We aligned the Xdrop-LRS data to the pBlueScript vector and found reads mapping to sequences encoding the f1 ori, the ampicillin resistance gene and the second ori (**Fig. S2D**). This part of the pBlueScript vector serves as backbone in many commonly used plasmids. It is therefore not surprising that vector sequences were in part identical. Since we did not detect any reads mapping to the pBlueScript-specific region located between the f1 ori and the second ori, we concluded that the vector sequence that integrated into the HepG2 target region originated from the CRISPR/Cas9 and not from the pBlueScript vector.

In sum, in the HepG2 Δt15 deletion clone, we observed a duplication of the target region, inversion of one copy and on-target integration together with sequences from the *E. coli* genome and the CRISPR/Cas9 vector.

### On-target genomic alterations occurred frequently

To estimate the approximate frequency of Cas9-induced genomic alterations, we tested whether on-target insertion events occurred in our other HAP1 and HepG2 Δt clones expected to carry the deletion. Since our Xdrop-LRS contig suggested an integration of genomic sequences larger than 7,900 bp, we could not quantify the frequency of on-target events by PCR using primers spanning the 5’ and 3’ flanking regions due to the very long amplicon length. We therefore designed a pair of primers annealing inside the target region (**Fig. S1E-F**). For HAP1, we inspected five HAP1 clones of which three contained the expected homozygous deletion because no target region-specific PCR product was detected (**Fig. 3D, Fig. S1C and E)**. The HAP1 Δt72 deletion clone showed two bands (∼340 bp and ∼1,200 bp) indicative for a target region duplication (**Fig. S1E**). In addition to the HAP1 Δt72 deletion clone, a PCR product with a length around 340 bp was detected in the HAP1 Δt19 deletion clone (**Fig. S1E**), suggesting a potential on-target genomic alteration.

In HepG2 cells, we tested the HepG2 Δt15 and 16 additional deletion clones in which the target region was deleted (**Fig. S1D**). Although we had validated the deletion in these 16 HepG2 clones by PCR, we still detected the target region in seven deletion clones (**Fig. 3D, Fig. S1F**). In sum, aberrant genomic changes at the on-target locus occurred frequently in HAP1 (40%) and HepG2 (47%) validated deletion clones (**Fig. 3D**).

### DNA fragments deriving from the CRISPR/Cas9 vector but also from other chromosomes can integrate at the inverted deleted target site in HepG2 deletion clone Δt8

Among the clones suspected to harbor on-target rearrangements detected by PCR, we further characterized the HepG2 Δt8 deletion clone by Xdrop-LRS. As in the HAP1 Δt72 and HepG2 Δt15 deletion clones, the alignment of Xdrop-LRS data to the human reference genome showed a decent coverage at the target locus **(Fig. S2B)** and a similar pattern of read discontinuity at DSB sites **(Fig. S2A)**. By applying our customized *de novo* assembly-based approach, we obtained two contigs.

Contig 1 contained four break points (BPs) **(Fig. 4A)**. BP1 connected the integrated CRISPR/Cas9 vector and an inverted target region-derived fragment, followed by inverted DNA fragments of 967 bp and 348 bp from chromosome 21 and 8 (BP2 and BP3), respectively. BP4 linked these fragments to the 3’ flanking region on chromosome 17. The fragment of chromosome 21 was derived from the gene body of lincRNA TCONS_I2_00017505 and the fragment of chromosome 8 contained sequences mapping to truncated LINE1 element sequences. Contig 2 consisted of three break points **(Fig. 4B)**. BP2 connected the inverted target region to the CRISPR/Cas9 vector fragment. BP1 and BP3 linked them to the 5’ and 3’ flanking regions, respectively. Alignment of both raw and corrected Xdrop-LRS reads to these contigs supported the on-target altered genomic content in the HepG2 Δt8 deletion clone **(Fig. 4A-B)**. Detailed inspection of the integrated fragments of CRISPR/Cas9 vector revealed differences between two contigs. In contig 1, the vector-derived fragment contained DNA sequences ranging from the puromycin resistance gene to the ampicillin resistance gene. While in contig 2, DNA sequences ranging from the U6 promoter to the chicken ß-actin promoter were integrated. Similar to the HepG2 Δt15 deletion clone, we only found reads carrying the sequence of gRNA-Δt-1 but not gRNA-Δt-2 **(Fig. 4C**).

**Fig 4.**
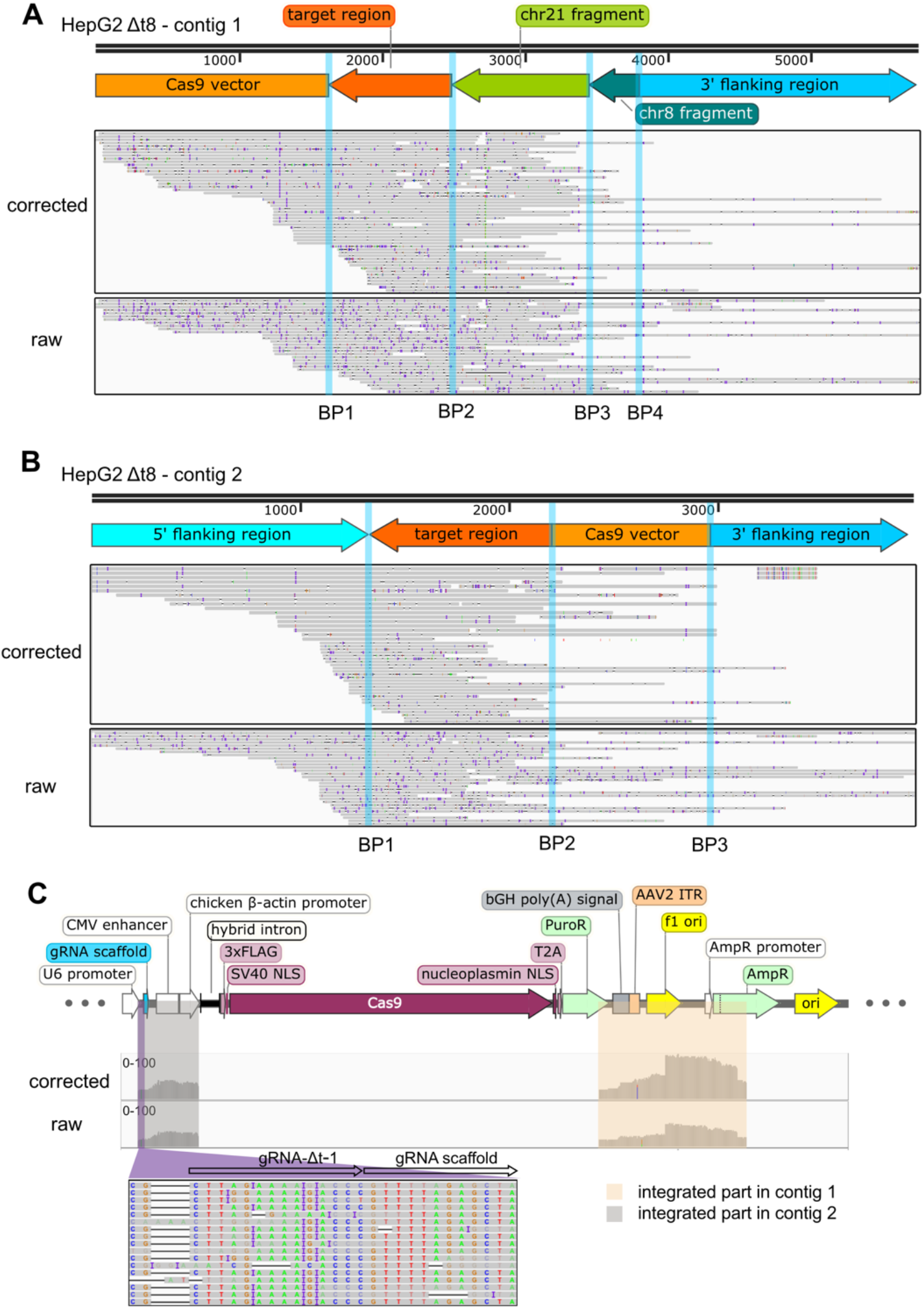
Clustered interchromosomal rearrangements, inversion of target-derived fragments and vector integration were identified in the HepG2 Δt8 deletion clone. **(A-B)** The hg38 genome browser view displays the Xdrop-LRS corrected (top) and raw (bottom) reads aligned to the *de novo* assembled contig 1 (A) and contig 2 (B) supporting different breakpoints (blue vertical lines) within the contig. Arrows (top) show genomic orientation and approximate size of the flanking (blue) and target (dark orange) regions as well as fragments deriving from the chromosome 21 (light green), the chromosome 8 (dark green) and the CRISPR/Cas9 transfection vector (light orange) present within the contig. **(C)** Coverage tracks show Xdrop-LRS corrected and raw reads aligning to the CRISPR/Cas9 vector (top). The vector fragments that were detectable in contig 1 (amber) and contig 2 (grey) are highlighted. The gRNA-Δt-1 cloning site and reads mapping to this region are highlighted (purple horizontal line) and magnified (bottom).

In sum, in the HepG2 Δt8 deletion clone, we observed an inversion of the target region and its on-target integration together with sequences from the chromosome 21 and 8 as well as the CRISPR/Cas9 vector.

### The occurrence of adverse on-target effects was independent of the sequence content of the deletion target region

To investigate whether the sequence similarity of the two tRNA-Cys-GCA genes caused the observed on-target genomic alterations, we repeated our dual gRNAs CRISPR/Cas9 approach in HAP1 and HepG2 cells using dual gRNAs that target the intergenic region (Δi) between our two target tRNA genes **(Fig. 5A-B)**. This intergenic region consists of unique DNA sequences without any gene annotation. Its deletion still allows transcription of the neighboring tRNA genes since the sequence motifs (A- and B-box) recognized by the Pol III promoter (A- and B-box) are located within the tRNA gene body (15, 16). For each Δi clone generated, we used primers flanking the intergenic region to validate a successful deletion event and internal primers to inspect potential genomic alterations **(Fig. 5B)**. We validated 7 HAP1 and 8 HepG2 Δi deletion clones, of those 1 (14%) HAP1 and 4 (50%) HepG2 Δi deletion clones carried the genomic alterations **(Fig. 5C, Fig. S3)**. Based on this frequency, we concluded that the sequence content of the deleted region did not cause the observed adverse on-target effects.

**Fig 5.**
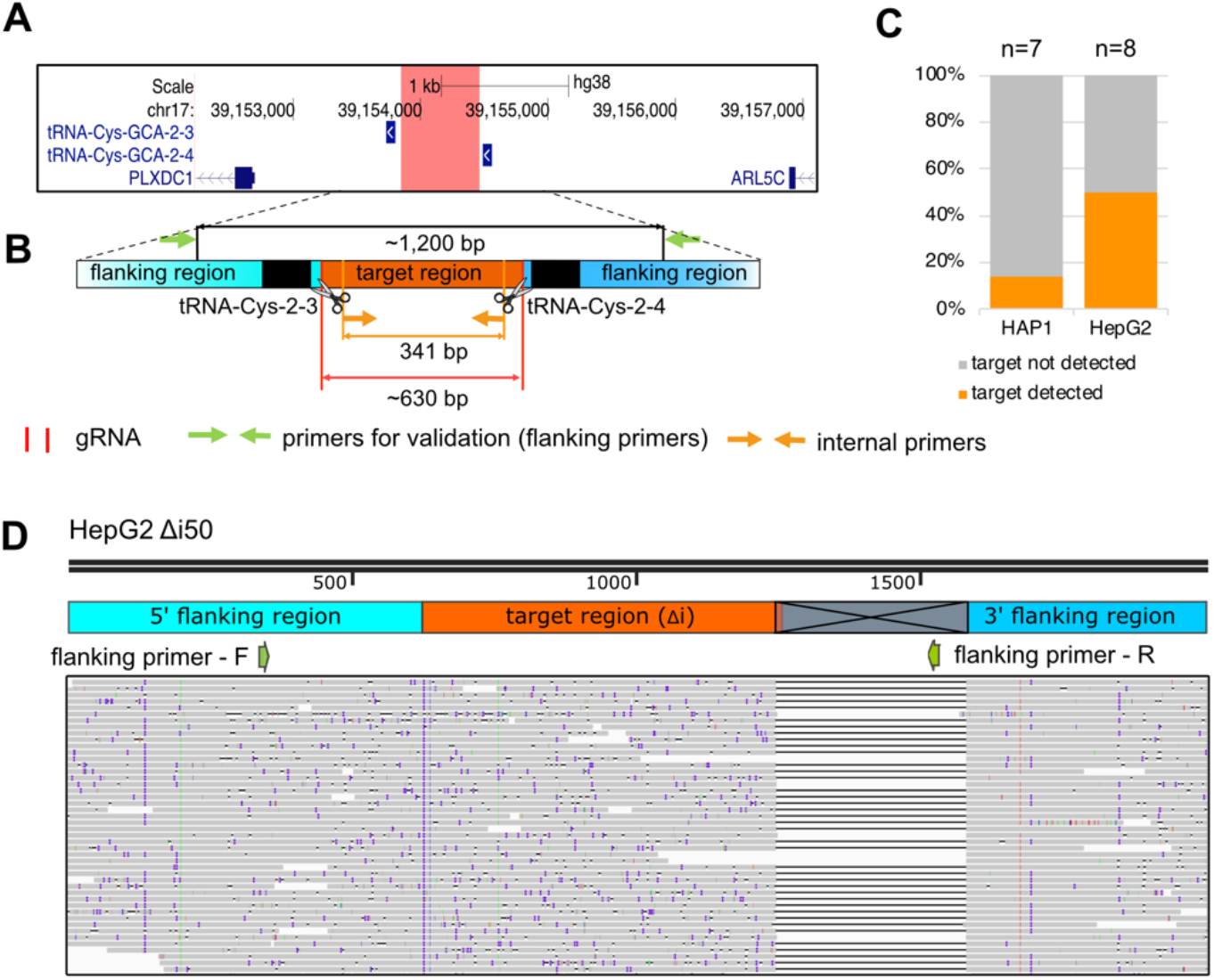
On-target genomic rearrangements were detected in intergenic region deletion clones. **(A)** The hg38 genome browser view shows the genomic location of the target locus (red) for intergenic region deletion (Δi). Arrows denote directionality of gene transcription. **(B)** Illustration of the design strategy for generating Δi clones. The cut sites of Cas9 (red horizontal lines and scissors), primers for validating the deletion (flanking primers, green arrows) and for detecting clones with genomic alterations (internal primers, light orange arrows) are indicated. The target region (orange) between two target tRNA genes (black) is around 630 bp. The size of the PCR product with flanking or internal primers in control clones is around 1,200 bp or 341 bp, respectively. **(C)** Stacked bar plot indicates the frequency of on-target genomic alterations in the validated HAP1 (n=7) and HepG2 (n=8) Δi deletion clones. **(D)** The hg38 genome browser view displays the alignment of the Xdrop-LRS raw reads to the human reference genome (hg38) at the target locus. The large deletion is indicated in the grey crossed box. The annealing sites of the flanking primers for the validation PCR experiment are displayed with green arrows.

To better understand the on-target rearrangements, we performed Xdrop-LRS on the HepG2 Δi50 deletion clone **(Fig. S3B)** by sequence enrichment of the deleted intergenic region **(Fig. S2B)**. We obtained high read coverage over the target region (**Fig. S2A**). The alignment of the raw reads to the human reference genome suggested a 331 bp deletion at the 3’ end of the target region **(Fig. 5D)**. This large deletion removed the binding site of the reverse primer used for identifying deletion clones, leading to a failure in detecting this genomic alteration by PCR. Notably, the deletion was located downstream of the second DSB site. Therefore, the target region itself remained still genomically intact.

### On-target genomic alterations led to functional DNA with biological consequences

We investigated potential biological consequences of the on-target genomic alterations by using several molecular and cellular assays.

First, we quantified tRNA gene usage across the different genotypes based on Pol III occupancy by ChIP-seq. Pol III ChIP-seq read counting is widely used for measuring tRNA gene usage (17–24). Unlike RNA-based methods, Pol III ChIP-seq provides tRNA gene-specific information and prevents inaccurate measurements caused by the complex tRNA structure and numerous modifications that block reverse transcription (61). After we normalized and applied a baseline cutoff, we counted Pol III ChIP-seq reads over the two target tRNA genes in the HAP1 Δt72 and HepG2 Δt15 deletion clones. Compared to the unmodified control clones, we found a 51% and 57% reduction in tRNA gene usage in the deletion clones HAP1 Δt72 and HepG2 Δt15, respectively **(Fig. 6A)**. This result confirms our previous observations that the deleted tRNA genes were integrated into the genomic on-target locus and can still be transcribed by Pol III. Although, the deleted region was duplicated in both the HAP1 Δt72 and HepG2 Δt15 deletion clones, we did not observe doubling of tRNA gene usage (**Fig. S1C-D**). Therefore, the different alleles (with and without a duplicated tRNA gene region) could counterbalance each other’s contributions to the tRNA-Cys-GCA transcript pool, which resulted in an intermediate transcriptional phenotype.

**Fig 6.**
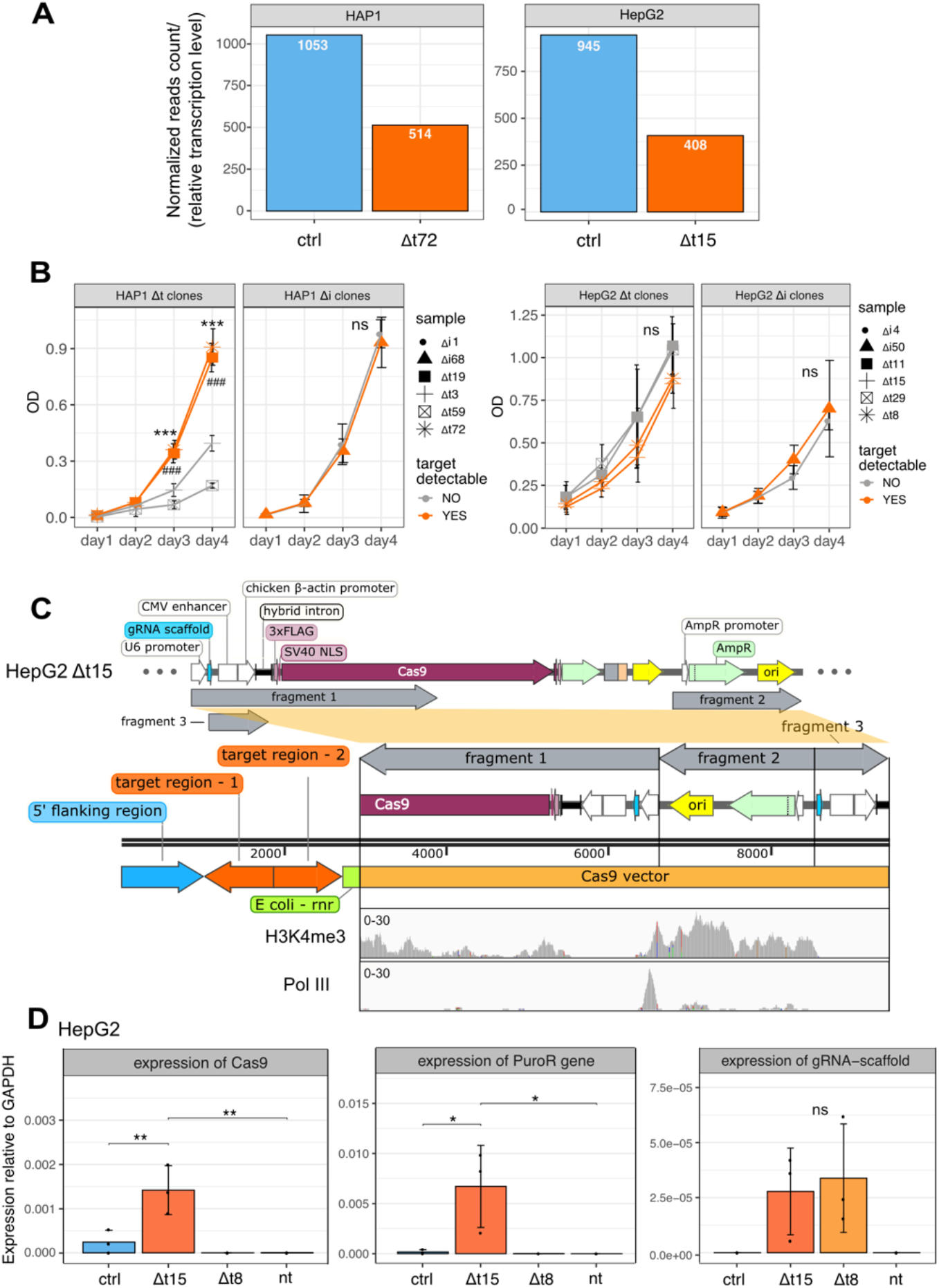
Adverse on-target genomic alteration affected cell growth and promoted active transcription. **(A)** Bar plots display the sum of Pol III ChIP-seq read counts covering the two target tRNA genes (tRNA-Cys-GCA-2-3 and tRNA-Cys-GCA-2-4) after normalization in the control clones and observed Δt (Δt72 and Δt15) deletion clones in HAP1 and HepG2 cell lines. The read counts are indicated at the top of the bar. **(B)** Line graphs show the relative number of cells (measured by optical density, OD) cultured over four days after seeding and measured by the crystal violet assay (n=3, mean +/- SD). Statistics: one-way ANOVA followed by Tukey HSD test. Significance code: p<0.001(***, when compared to the HAP1 Δt3 deletion clone and ###, when compared to the HAP1 Δt59 deletion clone); ns, not significant. **(C)** Illustration of the different CRISPR/Cas9 vector sequences (fragment 1-3, top) that integrated in the HepG2 Δt15 deletion clone (middle). The orientation of integration is shown. Coverage tracks (bottom) show mapping of H3K4me3 and Pol III ChIP-seq data to the integrated CRISPR/Cas9 vector sequences. **(D)** Bar graphs display expression levels of genes that integrated in the HepG2 Δt15 and Δt8 deletion clones. HepG2 non-targeting clones (ctrl) as well as non-transfected (nt) cells were used as controls. Expression levels were normalized to *GAPDH* gene expression and determined by qPCR (n=3, mean +/- SD). Statistics: one-way ANOVA, followed by Tukey HSD test. Significance codes: *p<0.05, **p<0.01, ns: not significant.

Since we confirmed that the target region-derived fragments in the HAP1 Δt72 and HepG2 Δt15 deletion clones were transcriptionally active, we next studied the cellular responses. We measured cell proliferation in the HAP1 and HepG2 Δt and Δi deletion clones with and without a detectable genomic on-target effect by using the crystal violet assay **(Fig. 6B)**. The HAP1 Δt72 and Δt19 deletion clones with on-target genomic alterations grew significantly faster than the *bona fide* HAP1 deletion clones Δt3 and Δt59. In contrast, the cell proliferation rate was nearly identical in the HAP1 Δi as well as in the HepG2 Δt and HepG2 Δi deletion clones. The observation that some cell clones gained a proliferative advantage could explain why frequencies of detectable target regions varied in the HAP1 Δt (2/5) and Δi (1/7) but not in the HepG2 Δt (8/17) and Δi (4/8) deletion clones **(Fig. 3D, Fig. 5C)**.

Lastly, we inspected the impact of the integrated sequences of the CRISPR/Cas9 vector. In the HepG2 Δt15 deletion clone, we found that the CRISPR/Cas9 vector was fragmented **(Fig. 6C)**. The integrated vector corresponded to three fragments. The first fragment started from the U6 promoter to the middle of Cas9 gene, and integrated into the HepG2 genome in inverse orientation, the second started from the ampicillin resistance gene to the ori element and also integrated in inverse orientation, and the third was composed of the gRNA-Δt-1 scaffold, the CMV enhancer and the chicken ß-actin promoter and integrated in the original orientation **(Fig. 6C)**. It would be expected that these exogenous sequences get silenced by the cell through heterochromatization, as it has been shown for other foreign sequences (62–64). To assess this, we inspected our H3K4me3 and Pol III ChIP-seq data and found that the plasmid-derived sequences were actively used in our deletion clone (**Fig. 6C**, H3K4me3 ChIP-seq track). Additionally, Pol III-binding to the U6 promoter gave rise to gRNA-Δt-1 **(Fig. 6**, Pol III ChIP-seq track). To quantify the expression of the integrated components, we performed a qPCR in the HepG2 Δt15 and Δt8 deletion clones using HepG2 non-targeting and non-transfected cells as baseline and negative controls, respectively. We detected expression of the Cas9 and puromycin resistance gene as well as of the gRNA-Δt-1 and its scaffold sequence in the HepG2 Δt15 deletion clone. In the HepG2 Δt8 deletion clone, only the gRNA-Δt-1 and its scaffold sequence were expressed, since the Cas9 and puromycin resistance genes were not integrated into the HepG2 genome according to our Xdrop-LRS data **(Fig. 4C)**. Our qPCR results confirmed that the integrated exogenous fragments were not silenced but instead actively transcribed.

In conclusion, local genomic rearrangements can lead to unwanted activation of gene expression and changes in cellular behavior, which can compromise the reliability of cell growth-dependent readouts commonly used in large-scale Cas9 screening. Our data therefore underscore the necessity to investigate potential on-target effects of Cas9-mediated deletion.

## DISCUSSION

We confirmed genomic on-target alterations in HAP1 and HepG2 cell clones after CRISPR/Cas9 induced cleavage by applying Xdrop-based enrichment for our target region followed by long-read sequencing and *de novo* sequence assembly. Although our genomic region of interest was cleaved by Cas9, we showed that the genomic fragment deriving from the target region was not eliminated from the nucleus. Instead, the cleaved fragment was duplicated, inverted and locally inserted into the genomes of HAP1 and HepG2 cells. In addition, we detected an integration of exogenous and endogenous DNA fragments as well as a large deletion downstream of the target region.

Based on our results, we reasoned that the HAP1 Δt72 deletion clone was heterogenous for the on-target genomic alterations (**Fig. S4A**). HAP1 is a near-haploid cell line but the haploid state is unstable and cells can rapidly become diploid (65). Although the exact chronologic order of events that occurred in this cell clone cannot be reconstructed, the observed genotype can be explained if the HAP1 Δt72 deletion clone derived from a diploid HAP1 cell. We speculate that after transfection of the CRISPR/Cas9 vector, Cas9 and the two gRNAs were efficiently expressed in the mother cell. Cas9 was directed via the dual gRNAs to cut the target-derived fragment from each of the two alleles in the diploid HAP1 mother cell. If the resulting DSB is not repaired before cell division, the cleaved fragments could be distributed randomly (66). In the case of the HAP1 Δt72 deletion clone, one daughter cell (DC1) could have used NHEJ to repair the DSBs resulting in a deletion. In contrast, the second daughter cell (DC2) would have employed MMEJ leading to the ligation and inversion of the two fragments at the original cut site during the repair process (58, 67). We postulate that the orientation of the fragment insertion was stochastic and can occur in either tandem (as in the HAP1 Δt72 deletion clone) or divergent (as in the HepG2 Δt15 deletion clone) orientation (**Fig. S4A-B**). Cells arising from DC1 and DC2 formed a heterogenous cell population, which also explains why, despite the duplication, ChIP-seq signals of Pol III and H3K4me3 over the target region were lower in cells deriving from the HAP1 Δt72 deletion clone when compared to the control clone (**Fig. 1C**).

Similar to our model for the HAP1 deletion clone, we consider that the HepG2 Δt15 deletion clone was heterogenous (**Fig. S4B**). HepG2 has a hyperploid karyotype with trisomy of chromosome 17 (68). Therefore, we speculate that two of the target-derived fragments were inserted into one of the three alleles in one daughter cell. We found that a large part of the integrated fragments of the CRISPR/Cas9 transfection vector included the gRNA sequence targeting the genomic region of interest and active promoters. Interestingly, a fragment originating from the *E. coli* genome was also inserted into the genome of the HepG2 Δt15 deletion clone. Since we used *E. coli* to produce large amounts of plasmid DNA, it is possible that bacterial over-lysis during plasmid preparation caused a release of *E. coli* genomic DNA, which got fragmented during plasmid purification. Small *E. coli* fragments can still bind to the purification column that selectively captures DNA of smaller size and get co-eluted with the plasmid DNA. We tested for any DNA contamination in our purified plasmid preparation by agarose gel electrophoreses. Although we were unable to visualize any DNA indicative of contamination (**Fig. S2E**), we cannot rule out that very low amounts of fragmented *E. coli* genomic DNA remained undetected.

In the HepG2 Δt8 deletion clone, the two obtained contigs indicated different on-target events with integration of different CRISPR/Cas9 vector sequences **(Fig. 4C)**. The two alleles that represent these two contigs could either reside in the same daughter cell (DC2, scenario 1) or both daughter cells (DC1 and DC2, scenario 2) **(Fig. S4C)**. In both scenarios, at least one allele carries a genuine deletion and was detectable initially through PCR validation, whereas the two other alleles with the inverted and integrated target derived-fragments were not captured by PCR due to their increased amplicon length **(Fig. S1D)**. It is notable that endogenous fragments from other chromosomes were found at the target site. We ruled out that these rearrangements were caused by translocations induced by Cas9 off-targeting at different chromosomal regions (69) because our gRNAs had very high on-target sequence specificity, and no sequence similarity between the gRNA and the chromosomal break sites were found. It is therefore unlikely that our dual gRNAs direct Cas9 in an off-target manner to cleave short fragments from chromosome 21 and 8, which would subsequently get integrated at the on-target region on chromosome 17. Instead, we suspected that this contig was the result of chromothripsis. A recent study showed that the Cas9 sgRNA system can trigger chromothripsis (66). Chromothripsis, also known as chromosome shattering, refers to extensive rearrangements within one or several chromosomes. Upon chromothripsis, chromosomes are fragmented and then configured in a random order mediated by NHEJ (70, 71). Therefore, we reasoned that chromothripsis occurred in the HepG2 Δt8 deletion clone when using our dual gRNA-Cas9 system, which lead to interchromosomal rearrangements at the target site.

In the HepG2 Δi50 deletion clone, we detected a large genomic deletion. Such deletions are the consequence of the Cas9 systems and can occur at different genomic loci and in many cell lines with various frequency (69, 72–74). Since the size of the large deletion is atypical for the NHEJ repair machinery (75, 76), other repair mechanisms such as MMEJ could play a role (69, 77, 78). However, we did not find evidence for sequence microhomology at the deletion breakpoint junction necessary for MMEJ **(Fig. S4D)**. Our findings demonstrate the complexity of DNA damage repair mechanism after Cas9-induced DSBs and limitations of predicting and validating genomic editing outcomes.

Additionally, the adverse on-target effect identified in our deletion clones revealed remarkable biological consequences. The integrated tRNA target region and exogenous fragments were bound by Pol III and H3K4me3 and resulted in increased cell proliferation in HAP1 cells. This adverse effect can hamper Cas9 screening approaches which often rely on cell growth as a readout.

Latest CRISPR engineering efforts have been focused on preventing potential on- or off-target effects. For example, restricting Cas9-induced DSB to G1 cells could minimize the formation of micronuclei and chromothripsis (66). Furthermore, the observed on-target effects described in this study may be mitigated by using Cas9-directed proteasomal degradation. That would limit the residence time of the Cas9 protein at the cut site and therefore enable accessibility of the DNA repair machinery and faster DSB repair (79, 80). Alternatively, suppressing MMEJ pathway may also reduce the frequency of the on-target rearrangements.

Xdrop technology combined with *de novo* long-read sequence assembly revealed unexpected complex genomic alterations that accompanied CRISPR/Cas9 deletions. Other methods could detect global Cas9 editing outcomes (69, 81–83). However, only few of them can reveal sequence aberrations in greater detail and with sufficient sequence length. For example, an integration at the region of interest can be detected with commonly available methods but those methods cannot find several large size integrations that occurred at the same time. Additionally, without performing a *de novo* assembly approach, it remains challenging to correctly decipher the fragmentation pattern of integrated DNA sequences. This is relevant since these fragments could have biological consequences. The workflow we introduced here is an enabling tool to study on-target outcomes of Cas9 deletion in single cell-derived clones.

Our approach revealed that Cas9 can induce duplications, inversions and local insertions of target-derived functional fragments as well as integration of exogenous and endogenous DNA fragments, possibly triggered by chromothripsis. Although it has been reported that a genomic region of interest can be either duplicated (58) or inverted (67) when using the dual gRNA system, our study provided direct evidence that the combination of a duplication and an inversion, along with an integration of exogenous DNA fragments and clustered interchromosomal rearrangements, can occur at the same time. Furthermore, we also demonstrated for the first time that despite these alterations, the target-derived fragments were still functional and can confound mechanistic interpretations. In conclusion, these findings present a new instance of unintended CRISPR/Cas9 editing events that can be easily overlooked and profoundly affect conclusions drawn from experimental read-outs.

## MATERIAL AND METHODS

### Cell culture

HAP1 cells were obtained from Horizon Discovery and grown in Iscove Modified Dulbecco Medium (Hyclone) supplemented with 10% Fetal Bovine Serum (Hyclone) and 1% Penicillin-Streptomycin (Sigma). HepG2 cells were obtained from American Type Culture Collection (ATCC, Rockville, MD) and grown in Dulbecco Modified Eagle Medium (Sigma) supplemented with 10% FBS and 1% Penicillin-Streptomycin. Cells were cultured in T75 flasks at 37°C and 5% atmospheric CO2. HAP1 and HepG2 cells were propagated by splitting 1/10 every two days and 1/4 every three days, respectively. Upon splitting, after the medium was aspirated, cells were washed with phosphate buffered saline (Sigma) and detached with 2mL of a trypsin-EDTA solution (Sigma). Trypsin was subsequently inactivated with a minimum of 3-fold surplus of culture medium before a cell fraction was passaged. Both cell lines have a certified genotype.

### Plasmid construction

gRNAs were designed and assessed using the design tool from the Zhang lab (https://zlab.bio/guide-design-resources). Each gRNA (Table S1) was separately cloned into the two BbsI restriction sites of pSpCas9(BB)-2A-Puro (px459) (2).

### Transfection and generation of single cell-derived clones

To enhance transfection efficiency, the short-size pBlueScript vector was co-transfected with large-size CRISPR/Cas9 vectors (3). For generating HAP1 and HepG2 deletion clones, two CRISPR/Cas9 vectors (Cas9-gRNA-1 and -2) were used. For creating non-targeting control clones (ctrl), no gRNA sequence was cloned into the CRISPR/Cas9 vector. Non-transfected (nt) cells served as additional control. About 160,000 HAP1 cells per well were plated in a 6-well plate and on the next day transfected using TurboFectin 8.0 (OriGene) according to the manufacturer’s instructions. Roughly 100,000 HepG2 cells were electroporated using the NEON electroporation system (Invitrogen) as previously described (3). After 24 h (HAP1) to 48 h (HepG2), cells were then selected by adding 2 µg/mL puromycin to the cell culture medium for two days. Afterwards, cells were allowed to recover in normal medium. Single-cell clones were hand-picked, grown and expanded.

### Genomic DNA extraction

HAP1 and HepG2 cells were lysed in 400 µL lysis buffer (0.5% SDS, 0.1M NaCl, 0.05M EDTA, 0.01M Tris HCl, 200 µg/mL proteinase K) and incubated at 55°C overnight. After which 200 µL of 5M NaCl were added, and the samples were vortexed and incubated on ice for 10min. After centrifugation (15,000×g, 4°C, 10 min), 400 µL of the supernatant were transferred to a new tube, mixed with 800 µL of 100% EtOH and incubated on ice for at least 10 min. Genomic DNA was pelleted by centrifugation (18,000×g, 4°C, 15 min), subsequently washed once with 70% EtOH and resuspended in nuclease-free water.

### PCR for genotyping

PCR was performed with Taq polymerase PCR (New England Biolabs) according to the manufacturer’s instructions. Briefly, 100 to 1000 ng of genomic DNA were used as template DNA. The 25 µL PCR reaction consisted of 1× standard Taq reaction buffer, 200 µM dNTPs, 0.2 µM primers (Table S1), and 1U Taq DNA polymerase. After thorough mixing of the PCR reaction, the subsequent PCR was performed using an initial denaturation step of 5 min at 95°C followed by 35 cycles of 30 sec at 95°C, 30 sec at 55 to 60°C, 60 sec per 1 kb at 68°C and a final extension step of 5 min at 68°C in a thermocycler (Applied Biosystems) with preheated lids. All PCR reactions were held at 4°C. PCR products were assessed by gel electrophoresis on a 1.2% agarose gel.

### Primer design

Primers (Table S1) were designed using NCBI Primer-BLAST (37) with default parameters (https://www.ncbi.nlm.nih.gov/tools/primer-blast/index.cgi?LINK_LOC=BlastHome). To ensure primer sequence specificity, we chose “Genomes for selected organisms” and “*Homo Sapiens*”. Primers with the highest specificity and GC-content of 40 to 60% were selected.

### Sanger sequencing

The PCR band indicative for the homozygous deletion (320 bp) was excised from the agarose gel and purified using Zymoclean Gel DNA Recovery Kit (Zymo Research) according to the manufacturer’s instructions. The purified DNA (75-150 ng) and the flanking sequence-specific primers (Supplemental Table 1) were sent for Sanger sequencing (Eurofins, Mix2Seq).

### RNA extraction and DNase-treatment

Cells were harvested at 80% to 90% confluency in 6 cm dishes. Cells were put on ice and 700 µL Qiazol (QIAGEN) were directly added onto the cells. Cells were scraped, mixed properly with Qiazol and collected in 1.5 ml tubes (Eppendorf). Afterwards, 140 µL chloroform were added. The mixture was shaken for 30 sec and incubated for 2.5 min at room temperature, before centrifugation at 9,000×g for 5 min at 4°C. After phase separation by centrifugation, the upper aqueous phase was carefully transferred in a new tube and mixed with 1 volume isopropanol. The tubes were inverted 5 times, followed by 10 min incubation at room temperature. The RNA was pelleted by centrifugation at 9,000×g for 10 min at 4°C. The pellet was washed carefully once using 700 µL ice-cold 70 % ethanol. The RNA was resuspended in 30-50 µL nuclease-free water. RNA concentration and purity were determined by nanodrop (Nanodrop 2000c). Afterwards, 6 µg RNA were mixed with 5 µL 10×TurboDNase buffer (Invitrogen), 1 µL TurboDNase (Invitrogen), 1 µL RNase Inhibitor (RiboLock, Invitrogen), and water was added to a total volume of 50 µL and subsequently incubated at 37°C for 30 min. DNase-treated RNA was purified with the Zymo RNA Clean & Concentrator Kit (Zymo Research) and eluted in 13 µL nuclease-free water. The concentration was determined by nanodrop.

### cDNA synthesis and qPCR

5 µg of purified total RNA were mixed with 1 µL random primers (250 ng/µL) and 1 µL dNTP Mix (10 mM each, Thermo Scientific) to a final volume of 12 µL. The mixture was incubated in a thermocycler for 5 min at 65°C and then quickly chilled on ice. Afterwards, 1 µL RNase-Inhibitor (RiboLock, Invitrogen), 4 µL 5x First-Strand Buffer (Invitrogen) and 2 µL 0.1 MDTT (Invitrogen) were added. After incubation for 2 min at 25°C, 1 µL SuperScript II reverse transcriptase (Invitrogen) was added. Samples were incubated in the thermocycler for 10 min at 25°C, 50 min at 42°C, 15 min at 70°C and held at 4°C.

After cDNA synthesis, 0.4 µL of *E. coli* RNase H (2 units) were used for digestion and incubated at 37°C for 20 min. Zymo DNA Clean & Concentrator Kit (Zymo Research) was used to purify cDNA.

The cDNA for qPCR was diluted to 1 ng/µL with nuclease-free water. The qPCR was conducted in 384-well plate with 10 µL system, containing 5 µL SYBR-Green Mix (PowerUp), 1 µL of primer, 1 µL of template and 3 µL of nuclease-free water. The qPCR was run on Real-Time-PCR machine (QuantStudio 5). The following parameters were set: initial heating steps of 2 min at 50°C followed by 2 min at 95°C. 40 cycles of 15 sec at 95°C and 1 min at 60°C. Melting curve: 15 sec at 5°C, 1 min at 60°C with ramp rate 1.6°C/sec and 15 sec at 95°C with ramp rate 0.075°C/sec. Gene expression levels of each target were normalized to the expression of *GAPDH*. Three technical replicates were used for each sample per target, and three biological replicates were included to calculate the mean and standard derivation.

### Crystal Violet assay

About 1,000 HAP1 or 5,000 HepG2 cells were seeded into 96-well plates in six technical replicates. Cells were assayed daily from day 1 to day 4. For the measurement, the medium was aspirated and carefully washed with 1×PBS twice. For staining, 100 µL of 0.5 % crystal violet solution (1% aqueous solution, diluted with water) (Sigma-Aldrich) were added to each well. After 1 min incubation at room temperature, the staining solution was removed. Each well was carefully washed 4 to 6 times with water to completely remove the unbound staining solution. After the wash, 100 µL of 10% acetic acid diluted in water (Fisher Scientific, AR grade) were used to dissolve the stain. The mixture was transferred to a new 96-well plate, and the absorption at 570 nm was measured on a plate reader (Spectramax i3x, Molecular Devices). Wells without cells were included in the assay as blank control. Background values were subtracted from the obtained values of optical density (OD). Three biological replicates were included to calculate the mean and standard deviation.

### Chromatin immunoprecipitation followed by sequencing (ChIP-seq)

ChIP-seq experiments were performed as previously described (23). Briefly, HAP1 and HepG2 cells were fixed in 1% formaldehyde, lysed, sonicated and then incubated with Pol III antibodies recognizing antigen POLR3A (21) or H3K4me3 antibodies (05-1339, Millipore). Immunoprecipitated DNA was used to generate sequencing libraries using the Takara SMARTer ThruPLEX DNA-seq Kit according to the manufacture’s protocol. Library size distribution and quality were assessed by an Agilent Bioanalyzer instrument using high-sensitivity DNA chips. The KAPA-SYBR FAST qPCR kit (Roche) was used to quantify the libraries. The sequencing run was performed with the NextSeq 500/550 High Output v2 kit (Illumina) for 81 cycles, single-end on an Illumina Nextseq500 platform.

### ChIP-seq data analysis

The qualities of reads were assessed using FastQC (38). Reads were aligned to the human reference genome hg38 using BWA (39). PCR duplicates and reads mapping to the ENCODE blacklist (https://sites.google.com/site/anshulkundaje/projects/blacklists) were removed using SAMtools (40) and NGSUtils (41). Bedgraph files were generated using deepTools (42). UCSC (http://genome.ucsc.edu) (43) or IGV (44) was used for visualization of the genomic loci or bam and bedgraph files. To avoid multi-mapping issues, we only considered reads with a mapping quality higher than 30 when inspecting peaks located at the target region. For aligning reads to the assembled contigs or the plasmids, BWA (39) was used for read mapping, followed by extraction of non-soft clipped reads with SAMtools (40). To count the number of Pol III reads, BEDTools (45) was used. Bed files containing tRNA gene location were obtained from GtRNAdb (46, 47).

### Xdrop

High-molecular-weight genomic DNA from the HAP1 Δt72 as well as HepG2 Δt15, Δt8 and Δi50 deletion clone was extracted using Quick-DNA kit (Zymo Research) and shipped to Samplix Services (Denmark) for Xdrop^®^ DNA enrichment followed by Oxford Nanopore Technology (ONT) long-read sequencing (LRS) (35, 48). Briefly, the Xdrop target enrichment workflow allows to capture and to enrich for a region of interest of up to 100 kb by targeting a short detection sequence. For the enrichment of the specific target region, detection primers were designed using the online primer design tool from Samplix to target a 120 bp detection sequence approximately mid-way between the Cas9 cut sites. At Samplix services, DNA quality was checked using the Tapestation™ System (Agilent Technologies Inc.) using Genomic DNA ScreenTape according to the manufacturer’s instructions. DNA was further purified using HighPrep™ PCR Clean-up Bead System according to the manufacturer’s instructions (MAGBIO Genomics) with the following changes: Bead-to-sample ratios were 1:1 (v:v) and elution was performed by heating the sample in the elution buffer for 3 min at 55°C before separation on the magnet. The samples were eluted in 20 µL 10 mM Tris-HCl (pH 8). Purified DNA samples were quantified by Quantus (Promega Inc.) Fluorometer™, according to the manufacturer’s instructions. PCR reagents, primers, as well as 4 -10 ng purified DNA of each sample, were partitioned in droplets and subjected to droplet PCR (dPCR) using the detection primers (see above). The dPCR droplets were then sorted by fluorescence-activated cell sorting (FACS). The isolated droplets were broken, and DNA was again partitioned in droplets and amplified by multiple displacement amplification in droplets (dMDA) reactions. After amplification, DNA was isolated and quantified. The MinION sequencing platform from ONT was used to generate LRS data from the dMDA samples as described by the manufacturer’s instructions (Premium whole genome amplification protocol (SQK-LSK109) with the Native Barcoding Expansion 1-12 (EXP-NBD104) and 13-24 (EXP-NBD114)). In short, 1.5µg amplified DNA of each sample was treated with T7 Endonuclease I, followed by size selection, end repair, barcoding, and adaptor ligation. After library generation, the samples were loaded onto a MinIon flow cell 9.4.1 (20fmol) and run for 16h under standard conditions as recommended by the manufacturer (Oxford Nanopore Inc.). Generated raw data (FAST5) was subjected to base-calling using Guppy v. 3.4.5 with high accuracy and quality filtering to generate FASTQ sequencing data.

### Xdrop data analysis

FASTQ reads were first corrected using Canu (49), followed by SACRA (50) to further identify and split chimeric reads. Reads were aligned to either the human reference genome (hg38) or the region of interest with minimap2 (51). For the *de novo* sequence assembly-based approach (**Fig. S2C**), the sequences of mapped reads were extracted from FASTQ files based on the mapped read IDs with SAMtools and SeqKit (52). The *de novo* sequence assembly was performed with these mapped reads using Canu or Raven (53). The output contigs were compared and assessed by pairwise alignment with Needle (https://www.ebi.ac.uk/Tools/psa/emboss_needle/), MegaBLAST against NCBI standard database of nucleotide collection (https://blast.ncbi.nlm.nih.gov/Blast.cgi) and manual inspection of the read alignment quality and coverage to the assembled contigs. If the flanking sequence of the target region needed to be extended, reads mapping to the 5’ or 3’ end of the contig were used for the second round of a *de novo* sequence assembly. If the read coverage was lower than the requirement of the assemblers, a manual extension was required. To do so, the soft clipped sequences of the reads mapping to the 5’ or 3’ end of the contig were visualized in the Integrative Genomics Viewer (IGV), manually compared and summarized. After the contig was successfully assembled, both corrected reads (using Canu and SACRA corrections) and raw reads were aligned to the contig. If there were more than one contig assembled for one sample, the contigs were merged into one *de novo* genomic reference to enhance the accuracy of the alignment. The contigs were assessed by visualization of aligned reads supporting breakpoints of the contig using IGV. ChIP-seq reads aligning to the contigs were used to assess and polish the contig.

## ACKNOWLEDGEMENTS

We are grateful for the fast and thorough delivery of enrichment data and support with the data analysis by the Samplix Services team, especially Christoffer Rozenfeld. We would like to thank the laboratories of Claudia Kutter, Marc Friedländer, and Vicent Pelechano, especially Eva Brinkman for critical discussions and feedback. We appreciated the technical support of Quim Perdices and Cristina Benito. We thank Kristoffer Sahlin, Philip Ewels and Remi-Andre Olsen for the helpful comments regarding our data analysis, Laura Baranello for the data interpretation and John Svetoft for the graphical design of our model. Our work benefited from the free usage of the SnapGene plasmid and contig viewer (https://www.snapgene.com/snapgene-viewer/) and free SMART Medical ART images (https://smart.servier.com). Computations were enabled by resources in project SNIC 2017/7-261, SNIC 2020/16-223, SNIC 2020/15-292, SNIC 2017/7-154 and uppstore2018110, provided by the Swedish National Infrastructure for Computing at UPPMAX.

## FUNDING

This work was supported by the Samplix Xdrop Grant Program (KG), Chinese Scholarship Council (201700260271 KG, CK), Knut & Alice Wallenberg foundation (KAW 2016.0174, CK), Ruth & Richard Julin foundation (2017–00358, 2018–00328, 2020-00294 CK), SFO-SciLifeLab fellowship (SFO_004, CK), Swedish Research Council (2019-05165, CK), Lillian Sagen & Curt Ericsson research foundation (2021-00427, CK), Gösta Milton’s research foundation (2021-00527, CK) and the Swedish National Infrastructure for Computing (SNIC) at UPPMAX.

## AUTHOR CONTRIBUTIONS

CK and KG conceptualized the project. KG, LGM, LW, AM, MS, YPS and JNS performed the laboratory experiments. KG did the analysis and visualized the data. RJW provided reagents. KG and CK acquired funding. KG and CK wrote the original draft. All authors contributed to the review and editing process.

## CONFLICT OF INTEREST

The authors declare no conflict of financial and non-financial interests.

## LIST OF ABBREVIATIONS

Δt: tRNA gene locus deletion
Δi: intergenic region deletion
BP: breakpoint
Cas9: CRISPR-associated protein 9
CRISPR: clustered regularly interspaced short palindromic repeats
ctrl: non-targeting control
DC: daughter cell
dMDA: droplet multiple displacement amplification
DSB: double-strand break
*E. coli*: *Escherichia coli*
gRNA: guide RNA
H3K4me3: histone 3 lysine 4 trimethylation
HAP1: chronic myeloid leukemia cell line
HepG2: hepatocellular carcinoma
LRS: long-read sequencing
MMEJ: microhomology-mediated end joining
NHEJ: nonhomologous end joining
nt: non-transfected control
ONT: Oxford Nanopore Technology
PAM: protospacer adjacent motif
Pol III: RNA polymerase III
tRNA: transfer RNA

## SUPPLEMENTAL MATERIAL

**Fig S1.**
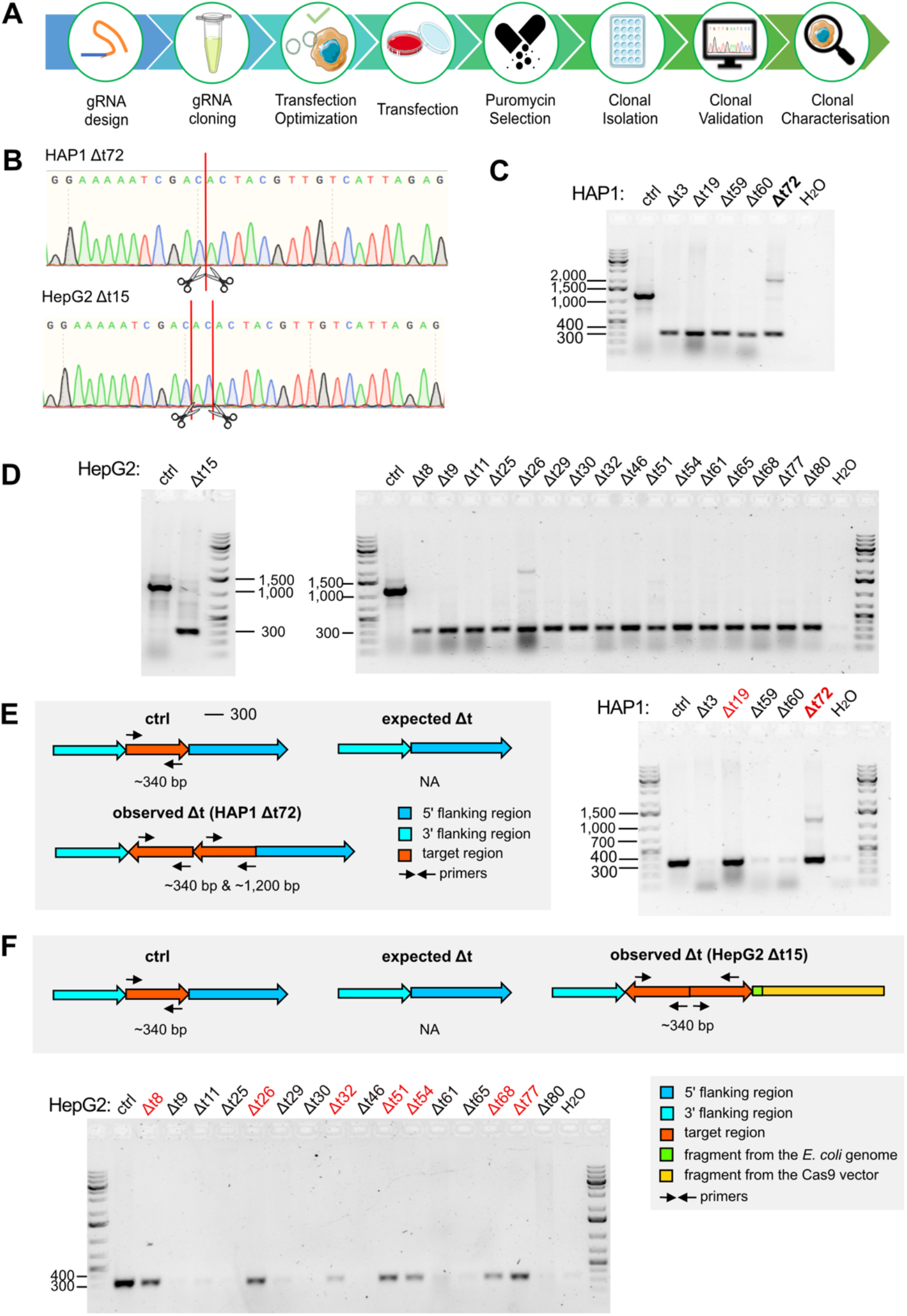
Validation of the tRNA gene locus deletion in HAP1 and HepG2. **(A)** Schematic illustration of the workflow to obtain single cell-derived deletion clones. **(B)** Chromatograms confirm deletions in the HAP1 Δt72 and HepG2 Δt15 deletion clones. DSB sites induced by CRISPR/Cas9 are shown by red lines and scissors. **(C-D)** Agarose gels confirm the size of the obtained PCR products to validate deletion events (320 bp) using primers annealing to the flanking regions of the target sites (Fig. 1B) in the HAP1 (C) and HepG2 (D) clones. In C, the HAP1 clone with duplicated target regions is bolded. (**E-F**) Schematic illustration of additional primers designed to distinguish unmodified control, expected and observed deletion events are used in HAP1 (E) and HepG2 (F). Deletion clones confirmed in C and D with a reinserted target region (340 bp, if duplication: 340 bp and 1,200 bp) are indicated in red. Marker bands specify DNA size in bp.

**Fig S2.**
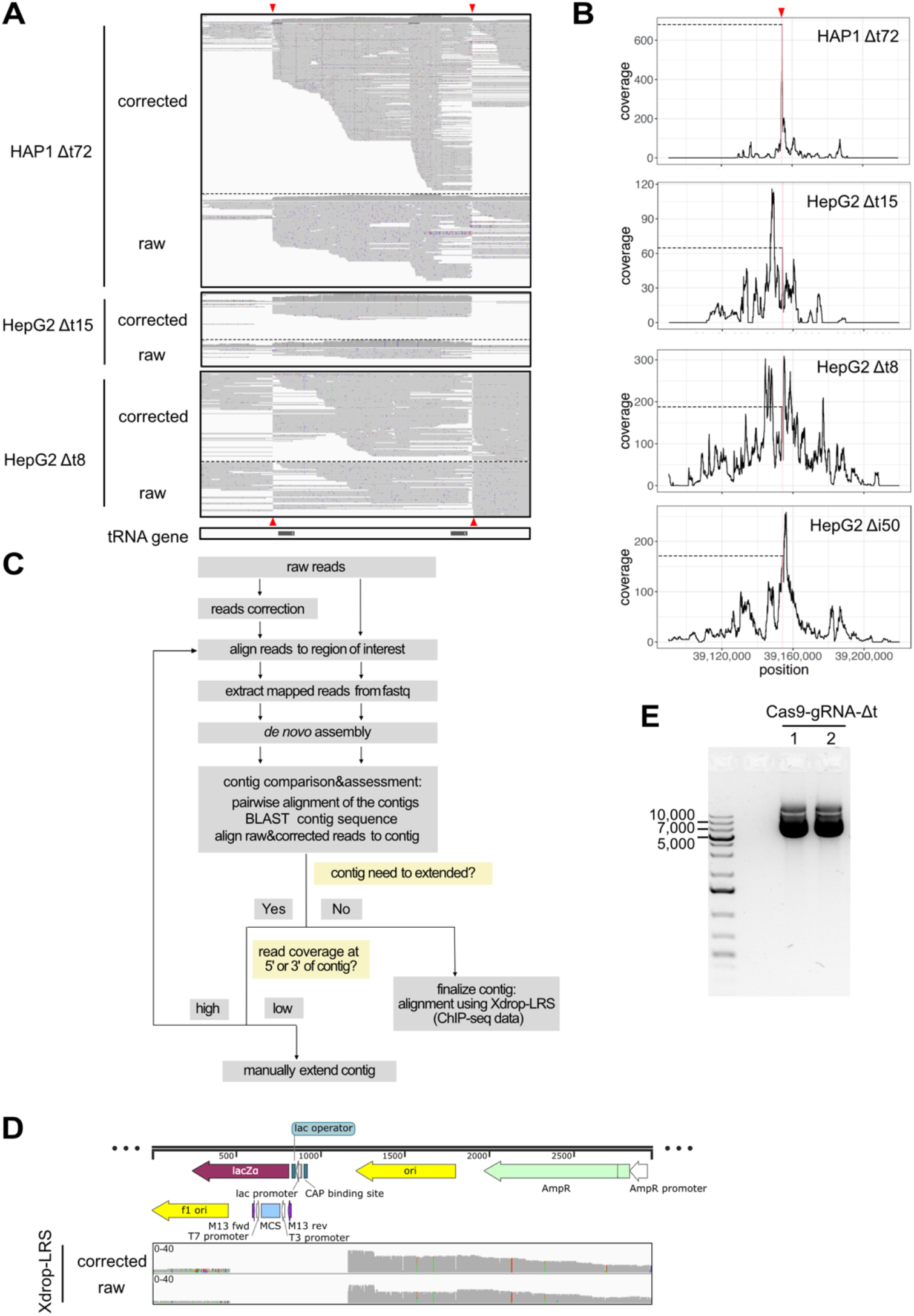
Complex on-target genomic alterations were revealed by Xdrop-LRS. (**A**) Alignment tracks display Xdrop-LRS corrected and raw reads at the target loci in HAP1 Δt72 (top), HepG2 Δt15 (middle) and HepG2 Δt8 (bottom) deletion clones. DSB sites (red triangles) and tRNA genes (black) are shown. (**B**) Line graphs show coverage when aligning raw reads obtained for the HAP1 Δt72 (top), HepG2 Δt15 (upper middle), HepG2 Δt8 (lower middle) and HepG2 Δi50 (bottom) deletion clones to the human reference genome (hg38). The position was defined by the coordinates from the human reference genome (hg38). Red lines and triangles highlight the location and dashed lines indicate the number of reads covering the target regions. (**C**) *De novo* assembly-based analysis workflow. (**D**) Coverage tracks display HepG2 Δt15 deletion clone Xdrop-LRS corrected and raw reads aligning to the pBlueScript vector. (**E**) Agarose gel confirms the size of the CRISPR/Cas9-gRNA-Δt-1 and -2 vectors (around 9,170 bp) after plasmid purification.

**Fig S3.**
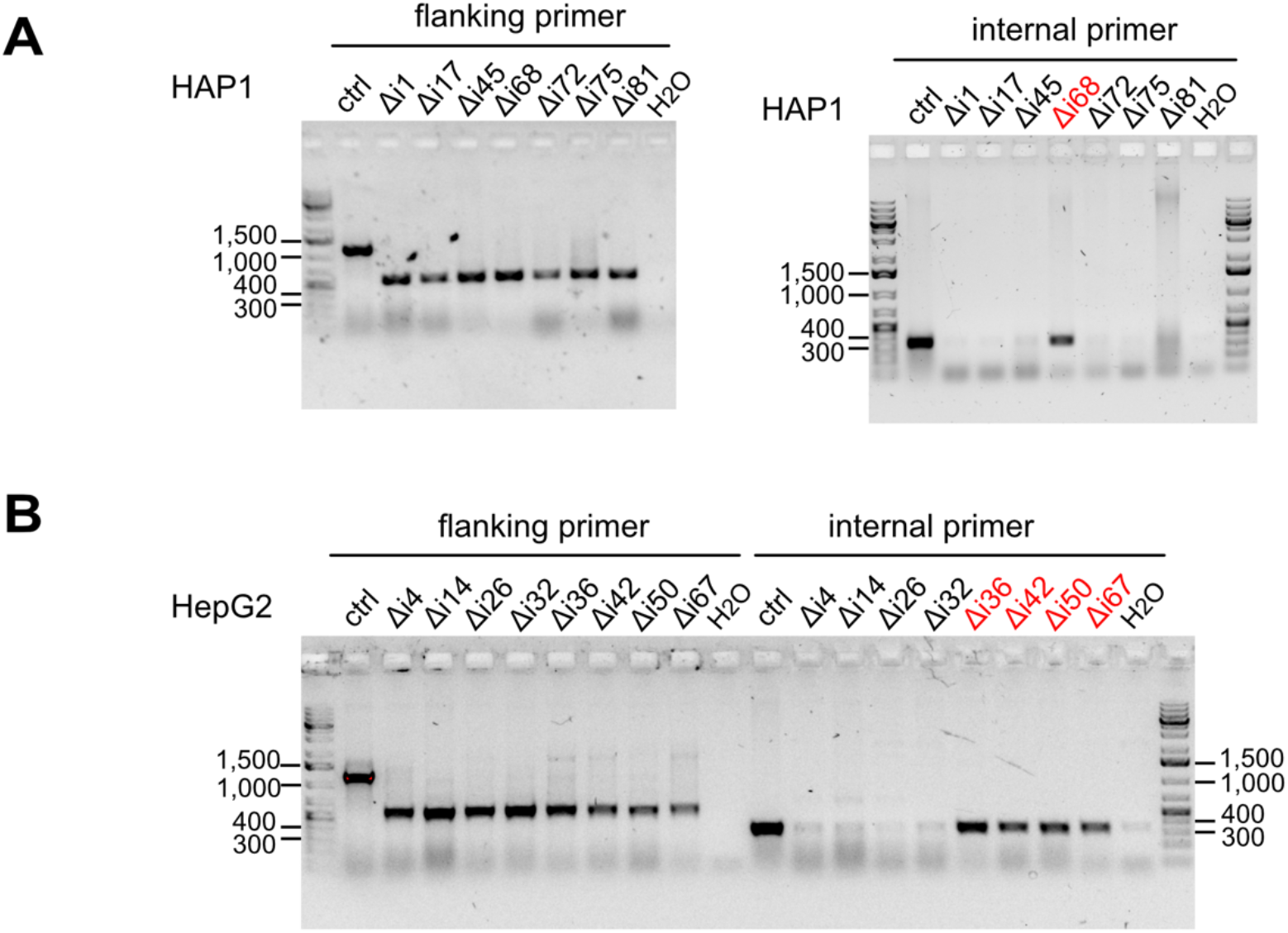
Validation of the intergenic region deletion in HAP1 and HepG2. **(A-B)** Agarose gels display the size of the obtained PCR products using flanking primer (left) to validate deletion events and the internal primer (right) to check for genomic alterations (Fig. 5B) in HAP1 (A) and HepG2 (B) clones. Clones with genomic alterations are indicated in red.

**Fig S4.**
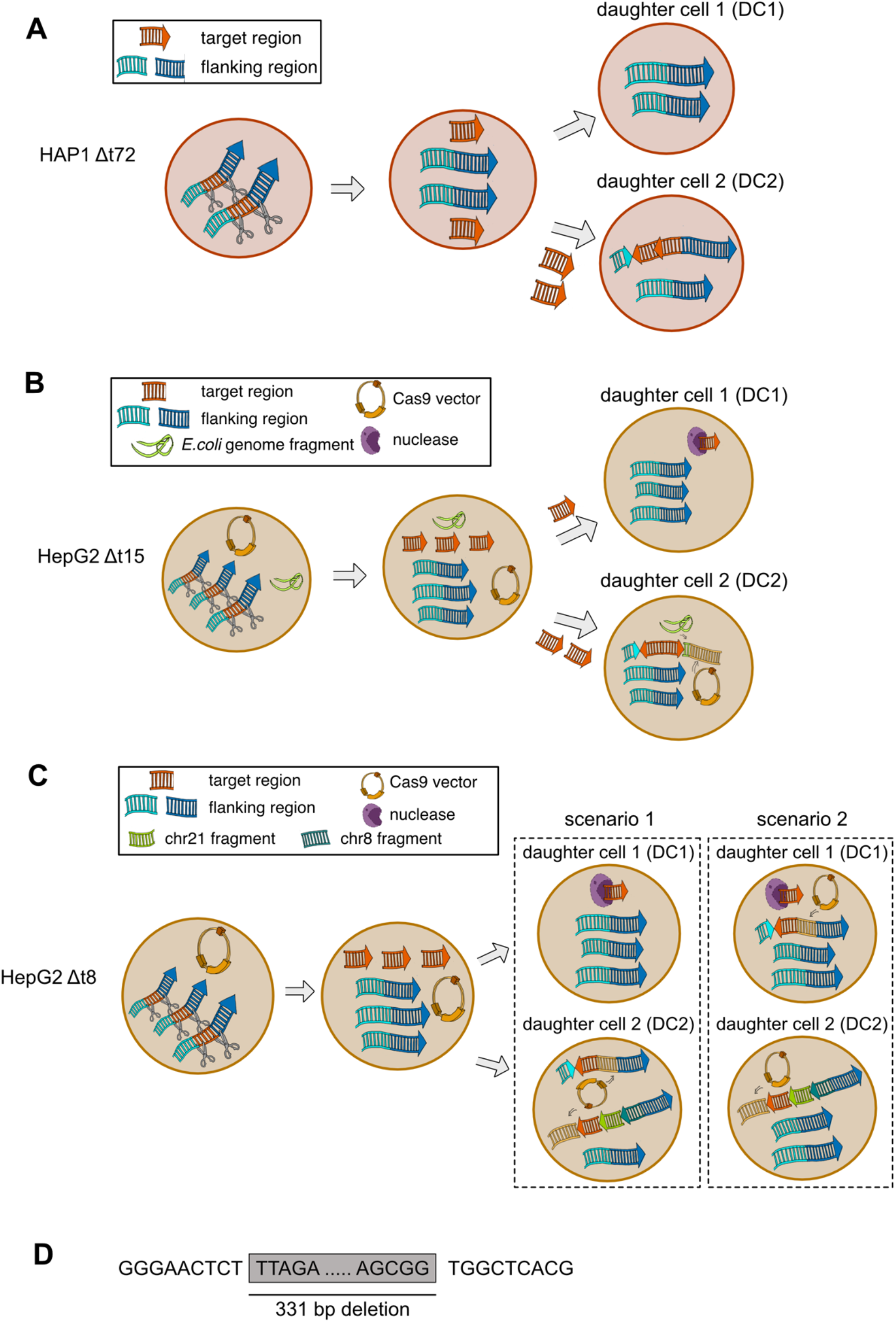
Hypothetical model of on-target genomic alterations in HAP1 and HepG2. **(A-C)** CRISPR/Cas9-mediated DSBs cleaving of the target region from the genome in the mother cell. Fragments were inverted and reinserted in one daughter cell (DC) in both the HAP1 (A) and the HepG2 (B-C) deletion clones. Additionally, the duplication (A-B) and the exogenous DNA sequences integration (B-C) as well as endogenous interchromosomal DNA integration (C) were observed. In the HepG2 Δt8 deletion clone (C), the two scenarios of the distribution of the two alleles with on-target genomic alterations are indicated with labels and dashed squares. (**D**) The sequence of the large deletion breakpoint junctions in the HepG2 Δi50 deletion clone is shown. The grey box indicates the deletion.

## Notes

### Competing Interest Statement

The authors have declared no competing interest.

## REFERENCES

1. Sander, J.D. and Joung, J.K. (2014) CRISPR-Cas systems for editing, regulating and targeting genomes. Nat. Biotechnol., 32, 347–350.

2. Ran, F.A., Hsu, P.D., Wright, J., Agarwala, V., Scott, D.A. and Zhang, F. (2013) Genome engineering using the CRISPR-Cas9 system. Nat. Protoc., 8, 2281–2308.

3. Søndergaard, J.N., Geng, K., Sommerauer, C., Atanasoai, I., Yin, X. and Kutter, C. (2020) Successful delivery of large-size CRISPR/Cas9 vectors in hard-to-transfect human cells using small plasmids. Commun. Biol., 3, 1–6.

4. Greene, E.C. (2016) DNA sequence alignment during homologous recombination. J. Biol. Chem., 291, 11572–11580.

5. McVey, M. and Lee, S.E. (2008) MMEJ repair of double-strand breaks (director’s cut): deleted sequences and alternative endings. Trends Genet., 24, 529–538.

6. Yang, H., Wang, H., Shivalila, C., Cheng, A., Cell L.S.- and 2013, undefined One-step generation of mice carrying reporter and conditional alleles by CRISPR/Cas-mediated genome engineering. Elsevier.

7. Maddalo, D., Manchado, E., Concepcion, C.P., Bonetti, C., Vidigal, J.A., Han, Y.C., Ogrodowski, P., Crippa, A., Rekhtman, N., Stanchina E. De, et al. (2014), In vivo engineering of oncogenic chromosomal rearrangements with the CRISPR/Cas9 system. Nature, 516, 423–428.

8. Canver, M.C., Bauer, D.E., Dass, A., Yien, Y.Y., Chung, J., Masuda, T., Maeda, T., Paw, B.H. and Orkin, S.H. (2014) Characterization of genomic deletion efficiency mediated by clustered regularly interspaced palindromic repeats (CRISPR)/cas9 nuclease system in mammalian cells. J. Biol. Chem., 289, 21312–21324.

9. Mansour, M.R., Abraham, B.J., Anders, L., Berezovskaya, A., Gutierrez, A., Durbin, A.D., Etchin, J., Lee, L., Sallan, S.E., Silverman, L.B., et al. (2014), An oncogenic super-enhancer formed through somatic mutation of a noncoding intergenic element. Science (80-.)., 346, 1373–1377.

10. Zhang, X., Choi, P.S., Francis, J.M., Imielinski, M., Watanabe, H., Cherniack, A.D. and Meyerson, M. (2016) Identification of focally amplified lineage-specific super-enhancers in human epithelial cancers. Nat. Genet., 48, 176–182.

11. Guo, Y., Perez, A.A., Hazelett, D.J., Coetzee, G.A., Rhie, S.K. and Farnham, P.J. (2018) CRISPR-mediated deletion of prostate cancer risk-associated CTCF loop anchors identifies repressive chromatin loops. Genome Biol., 19, 1–17.

12. Guo, Y., Xu, Q., Canzio, D., Shou, J., Li, J., Gorkin, D.U., Jung, I., Wu, H., Zhai, Y., Tang, Y., et al. (2015), CRISPR Inversion of CTCF Sites Alters Genome Topology and Enhancer/Promoter Function. Cell, 162, 900–910.

13. Antoniani, C., Meneghini, V., Lattanzi, A., Felix, T., Romano, O., Magrin, E., Weber, L., Pavani, G., Hoss S. El, Kurita, R., et al. (2018), Induction of fetal hemoglobin synthesis by CRISPR/Cas9-mediated editing of the human b-globin locus. Blood, 131, 1960– 1973.

14. Zhu, S., Li, W., Liu, J., Chen, C.H., Liao, Q., Xu, P., Xu, H., Xiao, T., Cao, Z., Peng, J., et al. (2016), Genome-scale deletion screening of human long non-coding RNAs using a paired-guide RNA CRISPR-Cas9 library. Nat. Biotechnol., 34, 1279–1286.

15. White, R.J. (2011) Transcription by RNA polymerase III: More complex than we thought. Nat. Rev. Genet., 12, 459–463.

16. White, R.J. (2005) RNA polymerases I and III, growth control and cancer. Nat. Rev. Mol. Cell Biol., 6, 69–78.

17. Canella, D., Bernasconi, D., Gilardi, F., LeMartelot, G., Migliavacca, E., Praz, V., Cousin, P., Delorenzi, M., Hernandez, N., Deplancke, B., et al. (2012), A multiplicity of factors contributes to selective RNA polymerase III occupancy of a subset of RNA polymerase III genes in mouse liver. Genome Res., 22, 666–680.

18. Oler, A.J., Alla, R.K., Roberts, D.N., Wong, A., Hollenhorst, P.C., Chandler, K.J., Cassiday, P.A., Nelson, C.A., Hagedorn, C.H., Graves, B.J., et al. (2010), Human RNA polymerase III transcriptomes and relationships to Pol II promoter chromatin and enhancer-binding factors. Nat. Struct. Mol. Biol., 17, 620–628.

19. Moqtaderi, Z., Wang, J., Raha, D., White, R.J., Snyder, M., Weng, Z. and Struhl, K. (2010) Genomic binding profiles of functionally distinct RNA polymerase III transcription complexes in human cells. Nat. Struct. Mol. Biol., 17, 635–640.

20. Barski, A., Chepelev, I., Liko, D., Cuddapah, S., Fleming, A.B., Birch, J., Cui, K., White, R.J. and Zhao, K. (2010) Pol II and its associated epigenetic marks are present at Pol III– transcribed noncoding RNA genes. Nat. Struct. Mol. Biol. 2010 175, 17, 629–634.

21. Kutter, C., Brown, G.D., Gonçalves, Å., Wilson, M.D., Watt, S., Brazma, A., White, R.J. and Odom, D.T. (2011) Pol III binding in six mammals shows conservation among amino acid isotypes despite divergence among tRNA genes. Nat. Genet., 43, 948–957.

22. Schmitt, B.M., Rudolph, K.L.M., Karagianni, P., Fonseca, N.A., White, R.J., Talianidis, I., Odom, D.T., Marioni, J.C. and Kutter, C. (2014) High-resolution mapping of transcriptional dynamics across tissue development reveals a stable mRNA-tRNA interface. Genome Res., 24, 1797–1807.

23. Rudolph, K.L.M., Schmitt, B.M., Villar, D., White, R.J., Marioni, J.C., Kutter, C. and Odom, D.T. (2016) Codon-Driven Translational Efficiency Is Stable across Diverse Mammalian Cell States. PLoS Genet., 12.

24. Gao, W., Gallardo-Dodd, C.J. and Kutter, C. (2021) Cell type-specific analysis by single-cell profiling identifies a stable mammalian tRNA-mRNA interface and increased translation efficiency in neurons. Genome Res., 10.1101/GR.275944.121.

25. Dieci, G., Fiorino, G., Castelnuovo, M., Teichmann, M. and Pagano, A. (2007) The expanding RNA polymerase III transcriptome. Trends Genet., 10.1016/j.tig.2007.09.001.

26. Ottenburghs, J., Geng, K., Suh, A. and Kutter, C. (2021) Genome Size Reduction and Transposon Activity Impact tRNA Gene Diversity While Ensuring Translational Stability in Birds. Genome Biol. Evol., 13.

27. Yeganeh, M. and Hernandez, N. (2020) RNA polymerase III transcription as a disease factor. Genes Dev., 34, 865–882.

28. Ishimura, R., Nagy, G., Dotu, I., Zhou, H., Yang, X.L., Schimmel, P., Senju, S., Nishimura, Y., Chuang, J.H. and Ackerman, S.L. (2014) Ribosome stalling induced by mutation of a CNS-specific tRNA causes neurodegeneration. Science (80-.)., 345, 455– 459.

29. Gomez-Roman, N., Grandori, C., Eisenman, R.N. and White, R.J. (2003) Direct activation of RNA polymerase III transcription by c-Myc. Nature, 421, 290–294.

30. Gingold, H., Tehler, D., Christoffersen, N.R., Nielsen, M.M., Asmar, F., Kooistra, S.M., Christophersen, N.S., Christensen, L.L., Borre, M., Sørensen, K.D., et al. (2014), A Dual Program for Translation Regulation in Cellular Proliferation and Differentiation. Cell, 158, 1281–1292.

31. Goodarzi, H., Nguyen, H.C.B., Zhang, S., Dill, B.D., Molina, H. and Tavazoie, S.F. (2016) Modulated expression of specific tRNAs drives gene expression and cancer progression. Cell, 165, 1416–1427.

32. Blanco, S., Dietmann, S., Flores, J. V, Hussain, S., Kutter, C., Humphreys, P., Lukk, M., Lombard, P., Treps, L., Popis, M., et al. (2014), Aberrant methylation of t RNA s links cellular stress to neuro-developmental disorders. EMBO J., 33, 2020–2039.

33. Van Bortle, K. and Corces, V.G. (2012) tDNA insulators and the emerging role of TFIIIC in genome organization. Transcription, 3, 277.

34. Raab, J.R., Chiu, J., Zhu, J., Katzman, S., Kurukuti, S., Wade, P.A., Haussler, D. and Kamakaka, R.T. (2012) Human tRNA genes function as chromatin insulators. EMBO J., 10.1038/emboj.2011.406.

35. Madsen, E.B., Höijer, I., Kvist, T., Ameur, A. and Mikkelsen, M.J. (2020) Xdrop: Targeted sequencing of long DNA molecules from low input samples using droplet sorting. Hum. Mutat., 10.1002/humu.24063.

36. Mikheyev, A.S. and Tin, M.M.Y. (2014) A first look at the Oxford Nanopore MinION sequencer. Mol. Ecol. Resour., 14, 1097–1102.

37. J, Y., G, C., I, Z., I, C., S, R. and TL, M. (2012) Primer-BLAST: a tool to design target-specific primers for polymerase chain reaction. BMC Bioinformatics, 13, 134.

38. Andrews, S., Krueger, F., Seconds-Pichon, A., Biggins, F. and Wingett, S. (2015) FastQC. A quality control tool for high throughput sequence data. Babraham Bioinformatics. Babraham Inst., 1, 1.

39. Li, H. and Durbin, R. (2009) Fast and accurate short read alignment with Burrows-Wheeler transform. Bioinformatics, 25, 1754–1760.

40. Li, H., Handsaker, B., Wysoker, A., Fennell, T., Ruan, J., Homer, N., Marth, G., Abecasis, G. and Durbin, R. (2009) The Sequence Alignment/Map format and SAMtools. Bioinformatics, 25, 2078–2079.

41. Breese, M.R. and Liu, Y. (2013) NGSUtils: A software suite for analyzing and manipulating next-generation sequencing datasets. Bioinformatics, 29, 494–496.

42. Ramírez, F., Ryan, D.P., Grüning, B., Bhardwaj, V., Kilpert, F., Richter, A.S., Heyne, S., Dündar, F. and Manke, T. (2016) deepTools2: a next generation web server for deep-sequencing data analysis. Nucleic Acids Res., 44, W160–W165.

43. Kent, W.J., Sugnet, C.W., Furey, T.S., Roskin, K.M., Pringle, T.H., Zahler, A.M. and Haussler, a. D. (2002) The Human Genome Browser at UCSC. Genome Res., 12, 996– 1006.

44. Robinson, J.T., Thorvaldsdóttir, H., Winckler, W., Guttman, M., Lander, E.S., Getz, G. and Mesirov, J.P. (2011) Integrative genomics viewer. Nat. Biotechnol., 29, 24–26.

45. Quinlan, A.R. and Hall, I.M. (2010) BEDTools: a flexible suite of utilities for comparing genomic features. Bioinformatics, 26, 841–842.

46. Chan, P.P. and Lowe, T.M. (2009) GtRNAdb: a database of transfer RNA genes detected in genomic sequence. Nucleic Acids Res., 37, D93–D97.

47. Chan, P.P. and Lowe, T.M. (2016) GtRNAdb 2.0: an expanded database of transfer RNA genes identified in complete and draft genomes. Nucleic Acids Res., 44, D184–D189.

48. Blondal, T., Gamba, C., Møller Jagd, L., Su, L., Demirov, D., Guo, S., Johnston, C.M., Riising, E.M., Wu, X., Mikkelsen, M.J., et al. (2021), Verification of CRISPR editing and finding transgenic inserts by Xdrop indirect sequence capture followed by short- and long-read sequencing. Methods, 10.1016/j.ymeth.2021.02.003.

49. Koren, S., Walenz, B.P., Berlin, K., Miller, J.R., Bergman, N.H. and Phillippy, A.M. (2017) Canu: Scalable and accurate long-read assembly via adaptive κ-mer weighting and repeat separation. Genome Res., 27, 722–736.

50. Impact of Chimera-less Long Reads on Metagenomics of Human Gut Viromes Treated With Multiple Displacement Amplification (2020) 10.21203/RS.3.RS-58640/V1.

51. Li, H. (2018) Minimap2: Pairwise alignment for nucleotide sequences. Bioinformatics, 34, 3094–3100.

52. Shen, W., Le, S., Li, Y. and Hu, F. (2016) SeqKit: A cross-platform and ultrafast toolkit for FASTA/Q file manipulation. PLoS One, 11, e0163962.

53. Vaser, R. and Šikić, M. (2020) Raven: A de novo genome assembler for long reads. bioRxiv, 10.1101/2020.08.07.242461.

54. Mali, P., Yang, L., Esvelt, K.M., Aach, J., Guell, M., DiCarlo, J.E., Norville, J.E. and Church, G.M. (2013) RNA-guided human genome engineering via Cas9. Science (80-.)., 339, 823–826.

55. Sternberg, S.H., Redding, S., Jinek, M., Greene, E.C. and Doudna, J.A. (2014) DNA interrogation by the CRISPR RNA-guided endonuclease Cas9. Nature, 507, 62–67.

56. Bosch-Guiteras, N., Uroda, T., Guillen-Ramirez, H.A., Riedo, R., Gazdhar, A., Esposito, R., Pulido-Quetglas, C., Zimmer, Y., Medová, M. and Johnson, R. (2021) Enhancing CRISPR deletion via pharmacological delay of DNA-PKcs. genome.cshlp.org, 10.1101/gr.265736.120.

57. Binda, C.S., Klaver, B., Berkhout, B. and Das, A.T. (2020) CRISPR-Cas9 Dual-gRNA attack causes mutation, excision and inversion of the HIV-1 proviral DNA. Viruses, 12, 330.

58. Shou, J., Li, J., Liu, Y. and Wu, Q. (2018) Precise and Predictable CRISPR Chromosomal Rearrangements Reveal Principles of Cas9-Mediated Nucleotide Insertion. Mol. Cell, 71, 498-509.e4.

59. Kraft, K., Geuer, S., Will, A.J., Chan, W.L., Paliou, C., Borschiwer, M., Harabula, I., Wittler, L., Franke, M., Ibrahim, D.M., et al. (2015), Deletions, inversions, duplications: Engineering of structural variants using CRISPR/Cas in mice. Cell Rep., 10, 833–839.

60. Hård, J., Mold, J.E., Eisfeldt, J., Tellgren-Roth, C., Häggqvist, S., Bunikis, I., Contreras-Lopez, O., Chin, C.-S., Rubin, C.-J., Feuk, L., et al. (2021), Long-read whole genome analysis of human single cells. bioRxiv, 10.1101/2021.04.13.439527.

61. Pinkard, O., McFarland, S., Sweet, T. and Coller, J. (2020) Quantitative tRNA-sequencing uncovers metazoan tissue-specific tRNA regulation. Nat. Commun., 11.

62. Taniguchi, Y., Nosaka, K., Yasunaga, J.I., Maeda, M., Mueller, N., Okayama, A. and Matsuoka, M. (2005) Silencing of human T-cell leukemia virus type I gene transcription by epigenetic mechanisms. Retrovirology, 2, 1–16.

63. Karlin ’, S., Doerfler, W. and Cardon’, L.R. (1994) Why is CpG suppressed in the genomes of virtually all small eukaryotic viruses but not in those of large eukaryotic viruses? J. Virol., 68, 2889–2897.

64. Liu, F., Campagna, M., Qi, Y., Zhao, X., Guo, F., Xu, C., Li, S., Li, W., Block, T.M., Chang, J., et al. (2013), Alpha-Interferon Suppresses Hepadnavirus Transcription by Altering Epigenetic Modification of cccDNA Minichromosomes. PLOS Pathog., 9, e1003613.

65. Olbrich, T., Mayor-Ruiz, C., Vega-Sendino, M., Gomez, C., Ortega, S., Ruiz, S. and Fernandez-Capetillo, O. (2017) A p53-dependent response limits the viability of mammalian haploid cells. Proc. Natl. Acad. Sci. U. S. A., 114, 9367–9372.

66. Leibowitz, M.L., Papathanasiou, S., Doerfler, P.A., Blaine, L.J., Sun, L., Yao, Y., Zhang, C.Z., Weiss, M.J. and Pellman, D. (2021) Chromothripsis as an on-target consequence of CRISPR–Cas9 genome editing. Nat. Genet. 2021 536, 53, 895–905.

67. Li, J., Shou, J., Guo, Y., Tang, Y., Wu, Y., Jia, Z., Zhai, Y., Chen, Z., Xu, Q. and Wu, Q. (2015) Efficient inversions and duplications of mammalian regulatory DNA elements and gene clusters by CRISPR/Cas9. J. Mol. Cell Biol., 7, 284–298.

68. Zhou, B., Ho, S.S., Greer, S.U., Spies, N., Bell, J.M., Zhang, X., Zhu, X., Arthur, J.G., Byeon, S., Pattni, R., et al. (2019), Haplotype-resolved and integrated genome analysis of the cancer cell line HepG2. Nucleic Acids Res., 47, 3846–3861.

69. Liu, M., Zhang, W., Xin, C., Yin, J., Shang, Y., Ai, C., Li, J., Meng, F.L. and Hu, J. (2021) Global detection of DNA repair outcomes induced by CRISPR–Cas9. Nucleic Acids Res., 49, 8732–8742.

70. Stephens, P.J., Greenman, C.D., Fu, B., Yang, F., Bignell, G.R., Mudie, L.J., Pleasance, E.D., Lau, K.W., Beare, D., Stebbings, L.A., et al. (2011), Massive Genomic Rearrangement Acquired in a Single Catastrophic Event during Cancer Development. Cell, 144, 27–40.

71. Ly, P., Teitz, L.S., Kim, D.H., Shoshani, O., Skaletsky, H., Fachinetti, D., Page, D.C. and Cleveland, D.W. (2016) Selective Y centromere inactivation triggers chromosome shattering in micronuclei and repair by non-homologous end joining. Nat. Cell Biol. 2016 191, 19, 68–75.

72. Kosicki, M., Tomberg, K. and Bradley, A. (2018) Repair of double-strand breaks induced by CRISPR–Cas9 leads to large deletions and complex rearrangements. Nat. Biotechnol., 10.1038/nbt.4192.

73. Shin, H.Y., Wang, C., Lee, H.K., Yoo, K.H., Zeng, X., Kuhns, T., Yang, C.M., Mohr, T., Liu, C. and Hennighausen, L. (2017) CRISPR/Cas9 targeting events cause complex deletions and insertions at 17 sites in the mouse genome. Nat. Commun. 2017 81, 8, 1– 10.

74. Wen, W., Quan, Z.J., Li, S.A., Yang, Z.X., Fu, Y.W., Zhang, F., Li, G.H., Zhao, M., Yin M. Di, Xu, J., et al. (2021), Effective control of large deletions after double-strand breaks by homology-directed repair and dsODN insertion. Genome Biol., 22, 1–22.

75. Guo, T., Feng, Y.L., Xiao, J.J., Liu, Q., Sun, X.N., Xiang, J.F., Kong, N., Liu, S.C., Chen, G.Q., Wang, Y., et al. (2018), Harnessing accurate non-homologous end joining for efficient precise deletion in CRISPR/Cas9-mediated genome editing. Genome Biol., 19, 1–20.

76. van Overbeek, M., Capurso, D., Carter, M.M., Thompson, M.S., Frias, E., Russ, C., Reece-Hoyes, J.S., Nye, C., Gradia, S., Vidal, B., et al. (2016), DNA Repair Profiling Reveals Nonrandom Outcomes at Cas9-Mediated Breaks. Mol. Cell, 63, 633–646.

77. Roidos, P., Sungalee, S., Benfatto, S., Serçin, Ö., Stütz, A.M., Abdollahi, A., Mauer, J., Zenke, F.T., Korbel, J.O. and Mardin, B.R. (2020) A scalable CRISPR/Cas9-based fluorescent reporter assay to study DNA double-strand break repair choice. Nat. Commun. 2020 111, 11, 1–15.

78. Owens, D.D.G., Caulder, A., Frontera, V., Harman, J.R., Allan, A.J., Bucakci, A., Greder, L., Codner, G.F., Hublitz, P., McHugh, P.J., et al. (2019), Microhomologies are prevalent at Cas9-induced larger deletions. Nucleic Acids Res., 47, 7402–7417.

79. Richardson, C.D., Ray, G.J., DeWitt, M.A., Curie, G.L. and Corn, J.E. (2016) Enhancing homology-directed genome editing by catalytically active and inactive CRISPR-Cas9 using asymmetric donor DNA. Nat. Biotechnol., 34, 339–344.

80. Brinkman, E.K., Chen, T., de Haas, M., Holland, H.A., Akhtar, W. and van Steensel, B. (2018) Kinetics and Fidelity of the Repair of Cas9-Induced Double-Strand DNA Breaks. Mol. Cell, 70, 801-813.e6.

81. Frock, R.L., Hu, J., Meyers, R.M., Ho, Y.J., Kii, E. and Alt, F.W. (2014) Genome-wide detection of DNA double-stranded breaks induced by engineered nucleases. Nat. Biotechnol. 2014 332, 33, 179–186.

82. Yin, J., Liu, M., Liu, Y., Wu, J., Gan, T., Zhang, W., Li, Y., Zhou, Y. and Hu, J. (2019) Optimizing genome editing strategy by primer-extension-mediated sequencing. Cell Discov. 2019 51, 5, 1–11.

83. Turchiano, G., Andrieux, G., Klermund, J., Blattner, G., Pennucci, V., el Gaz, M., Monaco, G., Poddar, S., Mussolino, C., Cornu, T.I., et al. (2021), Quantitative evaluation of chromosomal rearrangements in gene-edited human stem cells by CAST-Seq. Cell Stem Cell, 28, 1136-1147.e5.

